# Inhibition of autophagy-lysosomal function exacerbates microglial and monocyte lipid metabolism reprograming and dysfunction after brain injury

**DOI:** 10.1101/2025.09.04.674092

**Authors:** Amir A Mehrabani-Tabari, Nivedita Hegdekar, Brian R Herb, Sazia Arefin Kachi, Chinmoy Sarkar, Sagarina Thapa, Dexter PH Nguyen, Yulemni Morel, Mehari M Weldemariam, Ludovic Muller, W Temple Andrews, Marcia Cortes-Gutierrez, Xiaoxuan Fan, Natarajan Ayithan, Olivia Pettyjohn-Robin, Sabrina Bustos, Lacey K Greer, Jessica T Gore, Maureen A Kane, Seth A Ament, Jace W Jones, Marta M Lipinski

## Abstract

CNS has an overall higher level of lipids than all tissues except adipose and contains up to 25% of total body cholesterol. Recent data demonstrate a complex crosstalk between lipid metabolism and inflammation, suggesting potential contribution of the lipid-rich brain environment to neuroinflammation. While recent data support the importance of brain lipid environment to inflammatory changes observed in age related chronic neurodegenerative diseases, *in vivo* interactions between lipid environment, lipid metabolism and neuroinflammation in acute brain disease and injury remain poorly understood. Here we utilize a mouse model of traumatic brain injury (TBI) to demonstrate that acute neurotrauma leads to widespread lipid metabolism reprograming in all microglial and brain associated and infiltrating monocyte populations. Additionally, we identify unique microglial and monocyte populations with higher degree of lipid metabolism reprograming and pronounced accumulation of neutral storage lipids, including cholesteryl esters and triglycerides. These lipids accumulate not only in lipid droplets but also in the microglial and monocyte lysosomes and are associated with lysosomal dysfunction and inhibition of autophagy after TBI. Our data indicate that lipid accumulation in these cells is the result of altered lipid handling rather than lipid synthesis and is triggered by phagocytosis of lipid-rich myelin debris generated after TBI. Finally, we use mice with autophagy defects in microglia and monocytes to demonstrate that further inhibition of autophagy leads to more pronounced lipid metabolism reprograming and exacerbated cellular lipid accumulation. Our data suggest a pathological feedback loop, where lipid phagocytosis causes inhibition of autophagy-lysosomal function, which in turn exacerbates cellular lipid retention, reprograming and inflammation.

## Introduction

Recent studies reveal a complex bi-directional crosstalk between lipid metabolism and inflammation. This includes reports of immune stimuli altering cellular lipid composition and conversely, cellular lipid storage and lipid droplet formation promoting inflammatory responses.^1,2^ Deciphering the mechanisms of these interactions will be particularly important for understanding how lipid environment affects physiological and pathological immune responses in tissues high in lipid, such as the adipose, atherosclerotic plaques and the central nervous system (CNS), including brain and spinal cord.

Because of prevalence of branched cells with high membrane-to-volume ratios and presence of cholesterol-rich myelin, CNS has overall higher level of lipids than all tissues except the adipose and contains up to 25% of total body cholesterol.^3,4^ Altered expression of lipid metabolism genes is commonly observed in many age-and disease-associated microglial populations.^5–7^ Lipid metabolism reprograming is particularly pronounced in lipid droplet associated microglia (LDAM) which accumulate in the aged and Alzheimer’s disease (AD) brain.^1,8^ LDAM are highly pathogenic, with properties including inhibition of phagocytosis and high levels of inflammation. ^1^ In the context of AD accumulation of microglial lipid droplets is triggered by exposure to amyloid β and exacerbated by the presence of the AD-associated *APOE4* allele.^9,10^ It is less clear how their formation is initiated in the absence of amyloid pathology. It is also not known whether microglial lipid metabolism reprograming and/or LDAM generation are part of acute neuroinflammatory responses following infection or injury.

Traumatic brain injury (TBI) is a leading cause of death and disability worldwide, and a significant predisposing factor to development of neurodegenerative diseases later in life.^11^ An important component of TBI is exacerbated and prolonged inflammation. It involves several cell types, including both microglia and peripheral macrophages/monocytes infiltrating the brain after injury-induced blood-brain-barrier compromise.^12^ The reasons for the disproportionately high pro-inflammatory response after TBI remain poorly understood. We recently demonstrated that autophagy, a lysosome-dependent catabolic pathway necessary for degradation and recycling of proteins, protein aggregates, organelles and other cellular components,^13–15^ is inhibited after TBI and that multiple cell types including activated microglia and infiltrating monocytes are affected.^16–18^ In addition to its well-documented role in proteostasis and organelle quality control, autophagy has been shown to regulate inflammatory responses, with high levels of autophagy generally associated with anti-inflammatory, and low levels with pro-inflammatory phenotypes.^19,20^ Consistently, our data demonstrated that inhibition of microglial and monocyte autophagy contributes to excessive and prolonged neuroinflammation after TBI.^16^ Since recent data indicate that autophagy also participates in lipid catabolism by targeting and degrading lipid droplets through lipophagy,^21,22^ its inhibition in immune cells suggests potential interaction between autophagy, inflammation and lipid metabolism in the TBI brain.

The mechanisms behind autophagy inhibition in microglia and monocytes after TBI remain unknown. As the resident phagocytic cells of the brain, microglia are responsible for clearing dead cells and cellular debris including lipids, which are all delivered to lysosomes for degradation.^23,24^ In peripheral phagocytic cells such as atherosclerotic plaque macrophages, phagocytosis of certain lipids, in particular cholesterol, can cause lysosomal dysfunction.^25^ Similarly, myelin phagocytosis contributes to lysosomal dysfunction in multiple sclerosis (MS). In both atherosclerosis and MS lipid phagocytosis is also associated with accumulation of storage lipids and inflammation.^26–28^ Since TBI leads to generation of abundant myelin debris we hypothesized that similar mechanism could contribute to both the inhibition of autophagy and lipid metabolism reprograming after TBI.

We used multi-omics approach to investigate interaction between lipid metabolism, autophagy-lysosomal function and inflammation in monocytes after TBI. Our lipidomic data demonstrated that neutral lipids like triglycerides and cholesteryl esters accumulated in the activated microglial and monocytes in the vicinity of TBI lesion. We used single cell RNA sequencing (scRNA-seq) to demonstrated that TBI leads to widespread microglial and monocyte lipid metabolism reprogramming, which preferentially affected lipid handling as opposed to lipid synthesis genes. Additionally, our data identified microglial and monocyte populations with more pronounced lipid metabolism reprograming and demonstrated that they correspond to the lipid accumulating cells observed in the TBI lesion. Lipid accumulation in these cells was triggered by phagocytosis of lipid-rich myelin debris generated after TBI and occurred not only in lipid droplets but also in lysosomes, leading to their dysfunction and inhibition of autophagy. Confirming interaction between autophagy, lipid metabolism and inflammation, lipid metabolism reprograming and cellular lipid accumulation after TBI were exacerbated in mice with autophagy defects in microglia and monocytes. Our data suggest that TBI triggers a pathological feedback loop, where lipid phagocytosis causes inhibition of autophagy-lysosomal function, which in turn exacerbates cellular lipid retention, reprograming and inflammation.

## Results

### TBI leads to accumulation of neutral lipids in microglia and infiltrating macrophages

We used desorption electro-spray ionization - mass spectrometry imaging (DESI-MSI) to assess overall extent of TBI-induced changes in brain lipid content and distribution. Wild type C57BL/6 12-week-old male mice were subjected to moderate controlled cortical impact (CCI), a well-characterized mouse model of TBI. Brains were harvested 3 days after injury or sham surgery, a time point corresponding to peak inflammation and monocytes infiltration in the CCI model.^29^ We observed significant changes in brain lipid distribution after TBI (Figure S1A-F). This included prominent accumulation of neutral lipids such as cholesteryl esters and triglycerides, which was most pronounced in the vicinity of the TBI lesion (Figure 1A and Figure S1C). Accumulation of neutral lipids in the TBI tissue suggested increase in the formation of lipid storage organelles, the lipid droplets.^30^ To confirm this, we immunostained brain sections with antibodies against lipid droplet coating protein perilipin-3 (PLIN3). We observed lipid droplet formation in IBA1^+^ monocytes starting on day1, with peak at day 3 and decline by day 7 after TBI (Figure 1B). Neutral lipid accumulation in monocytes was confirmed by staining with neutral lipid dye, BODIPY (Figure 1C). We used flow cytometry to determine whether the affected cells were resident brain microglia (CD11B^+^ CD45^int^) or infiltrating monocytes (CD11B^+^ CD45^hi^) (Figure S1G). Consistent with immunostaining results, we observed significantly higher number of cells accumulating BODIPY starting on day 1 and peaking on day 3 after TBI. While neutral lipid accumulation was observed in both resident microglia and infiltrating monocytes, it was significantly more pronounced in the infiltrating monocyte populations (Figure 1D).

**Figure 1.**
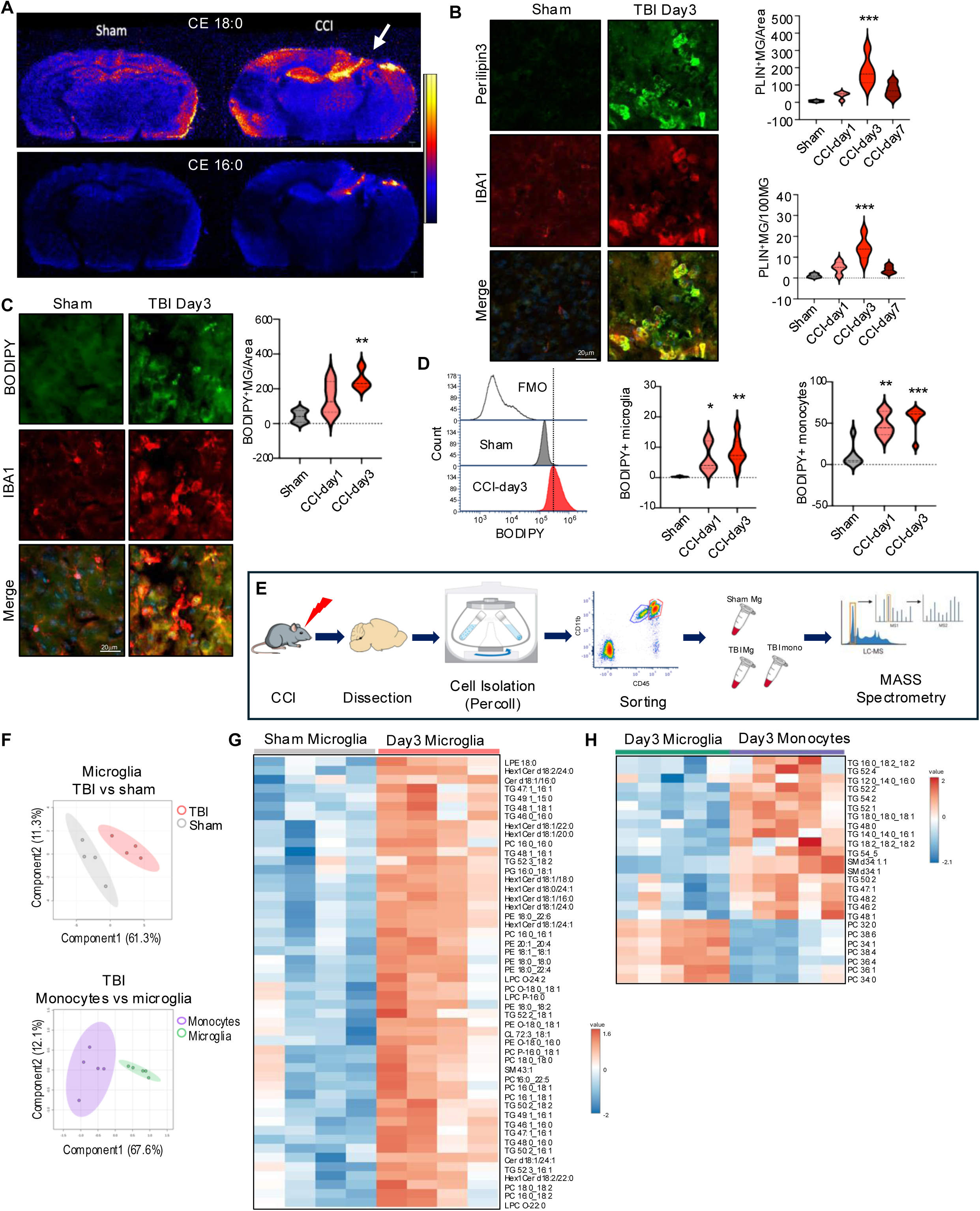
TBI leads to accumulation of neutral lipids in microglia and infiltrating macrophages. (A) Desorption Electrospray Ionization-Mass Spectrometry Imaging (DESI-MSI) of coronal mouse brain slices from sham and TBI (CCI model, day3), showing spatial distribution of cholesteryl esters 18:0 (cholesteryl stearate) and 16:0 (cholesteryl palmitate). (B) Immunostaining and quantification including time course of microglia/macrophages (IBA1, red) positive for lipid droplets protein, perilipin3 (green) in cortical sections from sham vs TBI mice (20X). Values are means ± SEM, n = 4 mice/group; *** P < 0.001 vs sham (one-way ANOVA with Tukey’s multiple comparisons test). (C) Immunostaining and quantification of neutral lipid (BODIPY, green) in microglia/macrophages (IBA1, red) in cortical sections from sham vs TBI. n = 4-5 mice/group; ** P < 0.01 vs sham (one-way ANOVA with Tukey’s). (D) Quantification of microglia and infiltrating macrophages accumulating neutral lipids (BODIPY) after TBI (day3). Left histogram demonstrating BODIPY signal shift from negative control (fluorescence minus one, FMO), in sham, and TBI microglia. Center & right quantification of BODIPY+ microglia and monocytes. n = 6-7 mice/group; * p>0.05, ** P<0.01, *** P<0.001 vs sham (one-way ANOVA with Tukey’s). (E) Experimental workflow for assessment of lipid composition of FACS isolated microglia and infiltrating monocytes from sham and TBI ipsilateral cortex. (F) Principal Component Analysis (PCA) scores plots demonstrating significant differences between lipid profiles of TBI vs sham microglia (top) and TBI monocytes vs TBI microglia (bottom) based on LC-MS/MS analyses. Each point represents sample form an individual mouse; 90% confidence intervals are shaded; n=4-5/group. (G-H) Heatmaps identifying differentially abundant lipids in TBI vs microglia (G) and TBI monocytes vs TBI microglia (H).

Our data suggested that following TBI mononuclear phagocytes including microglia and infiltrating macrophages accumulate neutral storage lipids such as triglycerides and cholesteryl esters. In cells other than adipocytes high level of storage lipid accumulation indicate overall lipid metabolism imbalance.^31^ To identify lipid signatures of these cells we used fluorescent activated cell sorting (FACS) to isolate microglia (CD11b^+^ CD45^int^) and infiltrating monocytes (CD11b^+^ CD45^hi^) from mouse perilesional cortex tissue (Figure 1E). Extracted lipids were analyzed using ultra-performance liquid chromatography (UPLC) coupled to selected reaction monitoring (SRM) on a tandem quadrupole mass spectrometer.^32^ We detected significant differences in lipid composition between microglia isolated from TBI versus sham mice as well as between infiltrating monocytes and microglia within the TBI group (Figure 1F). Consistent with immunostaining and flow cytometry results, the most prominent group of lipids with increased abundance in microglia after TBI were triglycerides, a storage lipid species highly abundant in lipid droplets (Figure 1G, Table S1). Triglyceride abundance was even higher in monocytes infiltrating the brain tissue after TBI (Figure 1H, Table S2). Other lipid species with increased abundance in TBI microglia and monocytes included hexosylceramides and sphingomyelins, lipids highly enriched in myelin,^33^ suggesting that internalized myelin debris from the TBI lesion could be a contributing source of accumulated lipid.

### TBI causes pronounced lipid metabolism reprogramming in microglia and macrophages

We performed single cell RNA sequencing (scRNA-seq) to gain a high-resolution picture of the changes in microglia and monocytes during the acute phase of TBI. We focused on day 3 after TBI, a period of rapid monocyte cell proliferation and inflammatory reprogramming.^34^ CD11B^+^ cells were isolated with magnetic beads from the ipsilateral hemispheres of TBI and sham mice (4/group) for 10x Genomics scRNA-seq, yielding 111,458 cellular transcriptomes after initial quality control (Figure 2A and Figure S2A). Cell clustering with Seurat revealed 29 transcriptionally distinct populations, which were annotated based on established marker genes and our experience with inflammatory changes in the CCI model^1,16,35–38^ (Figure 2B-C and Figure S2B). 22 out of 29 clusters (107,778 out of 111,458 sequenced cells, 96.7% enrichment) showed strong microglial/monocyte signatures expected for CD11B cells including 5 homeostatic microglia (Hom_MG_1-5), 5 surveillance microglia (Surv_MG_1-5), 5 disease associated microglia (DAM_1-5), and 7 brain associated macrophage (BAM_1-7) clusters. The remaining 7 minor clusters (each under 0.5% of total cell population) were excluded from further analysis.

**Figure 2.**
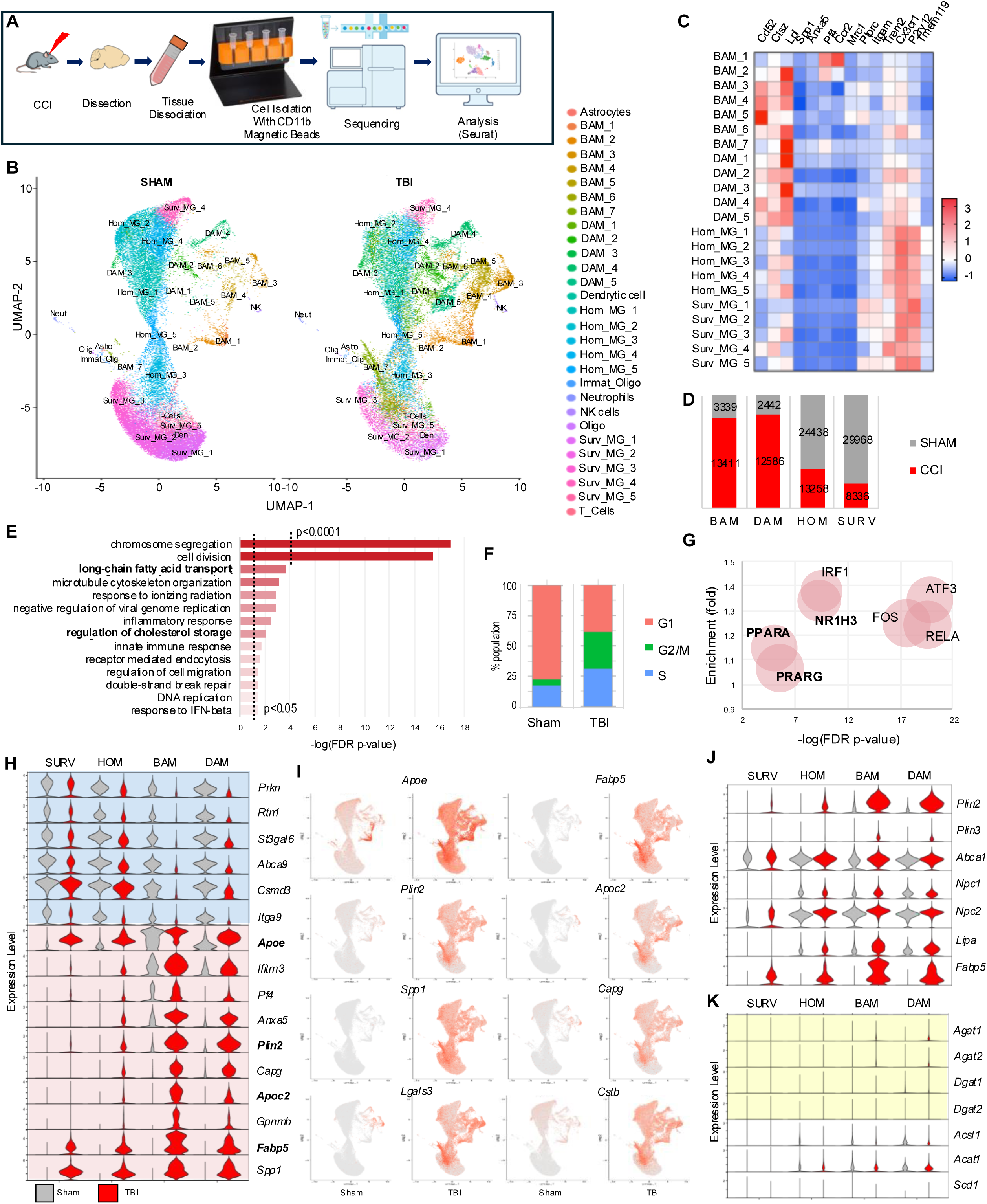
TBI causes pronounced reprogramming of lipid metabolism in microglia and macrophages. (A). Experimental workflow for assessment of transcriptomic changes in isolated CD11B^+^ cells from from sham vs TBI (day3) mouse brain. (B) Uniform manifold approximation (UMAP) demonstrating identification of 29 transcriptionally distinct clusters and their manually assigned identities. (C) Heatmap representing expression levels of canonical homeostatic and disease associated markers used for cluster identity assignment. Color coding is based on z-score scaling. (D) Bar chart showing distribution of the four main cell populations: surveillant microglia (SURV), homeostatic microglia (HOM), disease associated microglia (DAM), and brain associated macrophages (BAM) in sham vs TBI brain. (E) Gene Ontology Biological Process (GO:BP) enrichment analysis for genes differentially expressed in TBI vs sham. (F) Seurat Cell Cycle analysis demonstrating higher proportion of cells in S and G2/M phase of the cell cycle after TBI vs majority of cells in G1 phase in sham. (G) Bubble plot representing transcription factors link (TFlink) analysis. Identified as upstream regulators of DEGs in TBI vs sham. The size of each bubble is proportionate to the number of DEGs controlled by the transcription factor. (H) Violin plots demonstrating expression of top DEGs downregulated (blue) and upregulated (pink) in TBI in the four main microglial and macrophage cell populations. Highlighted in bold are lipid handling genes. (I) Feature plots demonstrating wide-spread changes in lipid metabolism (*Apoe*, *Fabp5*, *Plin2*, and *Apoc2*), microglial activation (*Spp1*), phagocytosis (*Capg*), and lysosomal fuction (*Lgals3* and *Cstb*) after TBI. (J-K) Violin plots demonstrating expression of lipid handling (J) and lipid synthesis (K) genes in the four main microglial and macrophage cell populations. Genes involved in triglyceride synthesis are highlighted in yellow in (K).

As expected, the majority of cells from sham mice belonged to homeostatic and surveillance microglial populations, with minor contribution of BAM and DAM. TBI lead to strong enrichment of the DAM and BAM populations, accompanied by a proportional decrease in homeostatic/surveillance microglia (Figure 2D). For initial analyses we compared 18318 differentially expressed genes (DEGs) in TBI and sham animals across all microglia/monocyte populations (Table S3). Gene Ontology Biological Process (GO:BP) enrichment analysis demonstrated overrepresentation of DEGs related to cell proliferation, which was confirmed by Seurat cell cycle analysis (Figure 2E-F, Figure S2C, Table S4). Other overrepresented categories included inflammatory responses, especially innate immunity known to be strongly activated after TBI, as well as terms related to lipid metabolism, especially fatty acid and cholesterol processing (Figure 2E). This was confirmed by transcription factor (TFlink) analysis, which in addition to canonical regulators of cell proliferation (FOS, ATF3) and inflammatory responses (RELA, IRF1) identified lipid metabolism regulators such as LXR (NRF1H3) and PPAR (PPARA, PPARG) as upstream drivers of the observed transcriptional changes (Figure 2G). These data suggest that TBI leads to general reprograming of microglial/monocyte lipid metabolism. To exclude the possibility that these results were driven by strong changes in a specific cell population, we compared expression of top up- and down-regulated genes in homeostatic and surveillance microglia, DAMs and BAMs from TBI versus sham mice. The top ten genes upregulated after TBI included four genes involved in lipid homeostasis (*Apoe*, *Fabp5*, *Plin2*, *Apoc2*), which were increased in most cell populations, with BAM and DAM affected to the highest extent, and surveillance microglia the least (Figure 2H-I, Figure S3A). Other genes dysregulated in multiple populations included those involved in phagocytosis (*Spp1*, *Capg*), lysosomal function (*Ctsb*) and lysosomal damage (*Lgals3*). Consistent with overall increase in DAM and BAM in TBI, many of these top markers are known DAM genes.

TLR4 signaling, which is strongly activated after TBI,^39^ has been shown to activate lipid synthesis leading to neutral lipid accumulation in TLR4 agonist treated mouse bone marrow derived macrophages (BMDM).^2^ To determine if this TLR4-dependent activation of lipid synthesis may be responsible for the observed accumulation of neutral lipid after TBI, we compared expression of genes involved in different lipid metabolism processes. Unexpectedly, while lipid handling genes were strongly upregulated in all cell types (Figure 2J), expression of lipid synthesis genes, including those involved in triglyceride synthesis, was affected to a much lesser degree (Figure 2K, Figure S3B). These data suggest that TBI-induced microglial and monocyte neutral lipid accumulation may be caused by altered cellular lipid handling rather than increased lipid synthesis. Finally, while our data identified transcription factors involved in regulation of lipid metabolism as some of the top upstream regulators responsible for the TBI-induced transcriptional changes, expression levels of mRNAs encoding these factors was not altered (Figure S3C).

### Lipid reprogramming is more pronounced in lipid-accumulating microglia and macrophages

While we observed TBI-induced lipid metabolism reprograming in all microglial and monocyte populations, some clusters were affected to a larger extent. In particular, BAM cluster 7 (BAM_7) and DAM cluster 3 (DAM_3) showed more pronounced upregulation of lipid handling genes as compared to other TBI cell populations (Figure 3A, Figure S3A). To determine whether these clusters may correspond to the lipid accumulating cells we observed after TBI (Figure 1), we used FACS to isolate microglial and monocyte population with high versus low levels of lipid accumulation (BODIPY^high^ and BODIPY^low^, respectively) from ipsilateral TBI day 3 cortices (Figure 3B). BODIPY^high^ microglia and monocytes showed elevated expression of BAM_7 and DAM_3 markers as compared to corresponding BODIPY^low^ cells (Figure 3C). Consistent with very low numbers of BODIPY positive cells in sham animals, DAM_3 and BAM_7 clusters were almost exclusively present in TBI samples. These data indicate that BAM_7 and DAM_3 represent the lipid-accumulating microglia and monocytes in the TBI brain.

**Figure 3.**
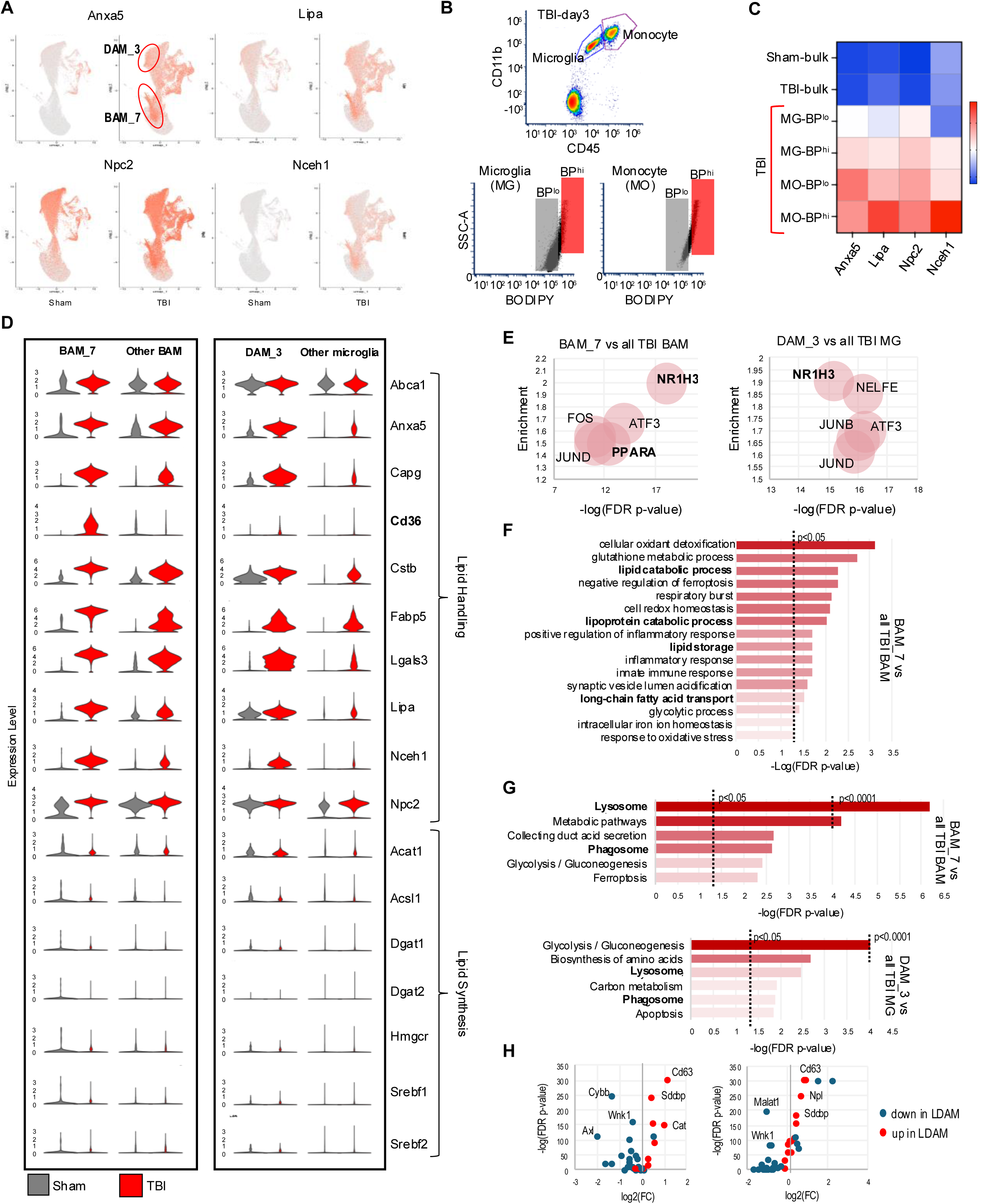
Lipid reprogramming is more pronounced in lipid-accumulating microglia and macrophages. (A) Feature plots for genes involved in lipid storage and processing (*Lipa*, *Npc2*, and *Nceh1*) and phagocytosis (Anxa5) identifying BAM_7 and DAM_3 as clusters with more pronounced lipid metabolism reporogramming. (B) Flow cytometry gating strategy for isolation of microglia (CD11B^+^CD45^int^) and monocytes (CD11B^+^CD45^hi^) with either low (BP^lo^) or high (BP^hi^) lipid load used for RNA isolation. (C) Heatmap of RTqPCR analysis of TBI-induced expression changes in DAM_3/BAM_7 markers from panel A in microglial (MG) and monocytes (MO) cells isolated using strategy from panel B. Increased expression levels in BPhi microglia and monocytes as compared to corresponding BPlo cells, indicated that that they represent DAM_3 and BAM_7 populations, respectively. (D) Violin plots representing expression levels of lipid handling genes (top) and lipid synthesis genes (bottom) in BAM_7 vs all other BAMs (left) and in DAM_3 vs all other microglia (right). (E) TFlink analysis identifying top five transcription factors responsible for DEGs in BAM_7 vs all other BAMs (left) and DAM_3 vs all other microglia (right). The size of each bubble is proportionate to the number of DEGs genes controlled by the transcription factor. (F) Gene Ontology Biological Process (GO:BP) enrichment analysis for genes differentially expressed in BAM_7 vs all other BAMs following exposure to TBI. Lipid metabolism terms are highlighted in bold. (G) KEGG pathway analyses of BAM_7 vs all other TBI BAMs and DAM_3 vs all other TBI microglia. Terms related to lysosomal function are highlighted in bold. (H) Volcano plots demonstrating altered expression of LDAM genes^1^ in BAM_7 vs all other TBI BAMs and DAM_3 vs all other TBI microglia.

To better understand the properties of lipid accumulating monocytes and microglia, we analyzed BAM_7 and DAM_3 clusters as compared to, respectively, all other BAMs and all other microglial cell populations. Both clusters demonstrated exacerbated induction of genes involved in lipid storage and handling and in lysosomal function after TBI (Figure 3D, Table S5). Notable were high expression levels of neutral cholesteryl ester hydrolase 1 (*Nceh1*), the enzyme responsible for converting neutral cholesterol esters to free cholesterol. We speculate that this could be a compensatory response to cholesteryl ester accumulation we observed in our DESI-MSI analyses (Figure 1A). We also noted almost exclusive upregulation of the scavenger receptor *Cd36* in BAM_7 after TBI which aligns with a recent report showing that *Cd36* positive macrophages in mouse model of ischemic stroke exhibited enhanced expression of phagocytic and lipid-handling genes.^40^ Taken together, these data suggest that high expression of *Cd36* may be a marker of lipid-loaded monocyte population after TBI.

Pathway analyses confirmed more pronounced dysregulation of lipid metabolism pathways in BAM_7 and DAM_3 as compared to other TBI BAM and microglial populations. This included identification of lipid metabolism transcription factors among the top 5 upstream regulators responsible for the observed transcriptional changes in both clusters (Figure 3E), and enrichment in processes related to cellular lipid catabolism, transport and storage in GO:BP analysis of BAM_7 (Figure 3F). These data are consistent with changes in lipid processing rather than synthesis as drivers of lipid accumulation after TBI. Other overrepresented processes included those involved in redox homeostasis, mitochondrial function and iron homeostasis, all previously reported to be affected after TBI. KEGG pathway analyses of both BAM_7 and DAM_3 included lysosomal and phagocytic pathways among the top six most significantly overrepresented, suggesting that in addition to changes in lipid metabolism, cells in both clusters may have altered lysosomal function as compared to other TBI BAM and microglial populations (Figure 3G). Lipid droplet associated microglia (LDAM) accumulate and play a pathological role during brain during aging and in neurodegenerative diseases such as AD.^1^ We compared expression of genes reported as most significantly up-and down-regulated in LDAM to those altered in BAM_7 and DAM_3. We observed overall strong correlation, with over 60% of LDAM genes significantly altered in both BAM_7 and DAM_3 (Figure 3H), suggesting that lipid accumulating microglia and monocytes acutely forming after injury share many features with those present in the aged and AD brain.

Recent data demonstrate that interferon signaling is an important contributor to TBI neuroinflammation.^34,41^ Consistently, we identified interferon regulatory factor IRF1 as one of the top upstream transcriptional regulators in TBI (Figure 2G). To investigate the relationship between lipid metabolism and interferon responses, we assessed expression of interferon response genes among identified cell populations. Our data demonstrate that unlike lipid metabolism, upregulation of interferon genes is restricted to specific BAM and DAM clusters (Figure S3D-E). Overall, BAM populations displayed more pronounced interferon responses as compared to DAM. However, we identified DAM_4 as displaying pattern of interferon gene expression similar to BAM; and DAM_2, which upregulated a unique set of interferon response genes, including *Ifit2* and *Ifit3*. Neither BAM_7 nor DAM_3 showed strong interferon responses as compared to other BAM and microglia, respectively. This suggests that lipid accumulation and upregulation of interferon pathways occur in different cell populations.

### Lipid accumulates in the lysosomes of microglia and macrophages after TBI

Our analyses of BAM_7 and DAM_3 indicated that both are characterized by changes in phagocytosis and lysosomal function. We investigated relationship between lipid accumulation and lysosomes by immunofluorescence for neutral lipid (BODIPY) and lysosomal membrane marker LAMP1 in IBA1^+^ microglial and monocyte cells (Figure 4A-B). We observed significant lysosomal accumulation of neutral lipid in phagocytes after TBI, which was further verified by 3D reconstruction of lipid-containing lysosomes (Figure 4C, Figure S4A). These data demonstrate that, in addition to lipid droplets, neutral lipids accumulate in the lysosomes rather than being processed for export to other cellular compartments, suggesting lysosomal dysfunction. To determine how TBI affects lysosomal composition and function, we isolated lysosome enriched fractions from perilesional cortical tissue at 1 and 3 days after TBI or sham surgery^17^ for LC-MS/MS lipidomic and proteomic analyses (Figure 4D). We detected significant differences in both lysosomal lipid and protein composition after TBI (Figure 4E). Lipidomic results demonstrated lysosomal accumulation of neutral lipids, including triglycerides and cholesteryl esters (Figure 4F, Figure S4B-C, Table S6). Significant enrichment for storage lipids normally observed in lipid droplets was confirmed by lipid ontology (LION) analysis (Figure 4G). Additionally, we observed increased abundance of ceramides, which together with cholesteryl esters suggested myelin as the source of lipids accumulating in TBI lysosomes (Figure 4F). Consistent with its pronounced transcriptional upregulation and its known role as the major lipid carrier in the brain,^42^ our proteomic data identified APOE as the top protein enriched in day 3 TBI lysosomes (Figure 4H, Table S7). Other increased proteins included ANXA5, a phagocytic marker transcriptionally elevated after TBI, and glycerol-3-phosphate dehydrogenase 1 (GPD1), an enzyme involved in triglyceride metabolism and the cause of transient infantile hypertriglyceridemia.^43^ Finally, we observed higher abundance of several members of the serpin family protease inhibitors (SERPINA1B, 1D, 1E, and 3K). Serpins irreversibly inhibit extracellular serine proteases involved in coagulation, cell migration and inflammatory responses^44^ and they entrapment in the lysosomes could contribute to TBI neuroinflammation.

**Figure 4.**
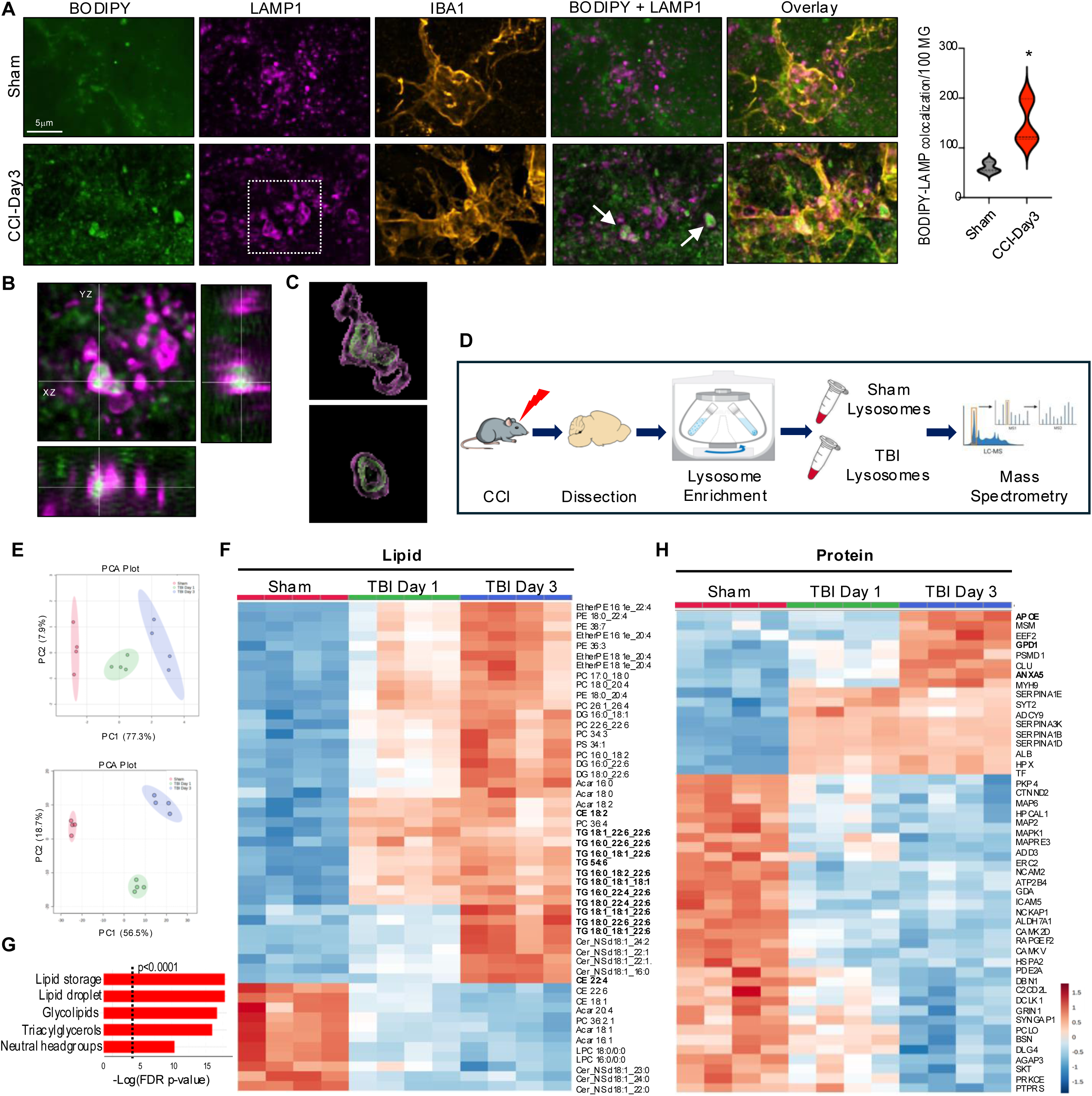
Neutral lipids accumulate in the lysosomes of microglia and macrophages after TBI. (A) Confocal images and quantification demonstrating accumulation of neutral lipid (BODIPY, green) in lysosomes (LAMP1, magenta) of microglia/macrophage cell (IBA1, orange) in coronal brain sections from TBI vs sham mice (100X); P < 0.05 using Student’s t-test. (B) Orthogonal views of indicated image area from panel A demonstrating localization of BODIPY within LAMP1 lysosomes. (C) 3D reconstruction of BODIPY engulfment inside lysosomes indicated with white arrow in panel A. (D) Experimental workflow for assessment of lipid and protein composition of lysosomal fractions from peri-lesion cortical tissue of TBI vs sham. (E) PCA plots demonstrating significant differences in lipid (top) and protein (bottom) composition between lysosomal fractions from sham vs TBI-day3 cortices based on LC-MS/MS analyses. Each point represents sample from an individual mouse; 90% confidence intervals are shaded; n = 4/group. (F) Heatmap identifying lipids differentially abundant in lysosomal fractions from TBI-day1 and TBI-day3 vs sham cortex based on ANOVA. (G) Lipid ontology (LION) enrichment analysis of differentially abundant lipids in TBI-day3 vs sham. (H) Metaboanalyst generated heatmap identifying proteins differentially abundant in lysosomal fractions from TBI-day1 and TBI-day3 vs sham cortex based on ANOVA.

### Myelin phagocytosis inhibits lysosomal function and autophagy after TBI

We recently demonstrated that autophagy is inhibited in both resident microglia and infiltrating monocytes after TBI, resulting in exacerbation of inflammatory responses.^16^ Both the timing (peak at day 3 post injury) and cell type distribution (higher in infiltrating monocytes than in microglia) of autophagy inhibition correlated with that observed for lipid accumulation after TBI (Figure 1). Since correlation between lipid accumulation and inhibition of autophagy flux in macrophages has been observed in the context of atherosclerosis,^45^ we tested whether autophagy flux is also inhibited in lipid accumulating monocytes of TBI lesion. Immunofluorescence staining showed that ∼72% of neutral lipid positive (BODIPY^+^) monocytes (IBA^+^) also accumulated high levels of autophagy adaptor p62, indicative of decreased autophagic cargo degradation (Figure 5A-B). Autophagy flux inhibition in lipid accumulating mononuclear phagocytes was verified using flow cytometry, where levels of both p62 and LC3 were significantly higher in BODIPY^+^ microglia and infiltrating monocytes in the TBI brain as compared to corresponding BODIPY^-^ cells (Figure 6C). Moreover, BODIPY^+^ populations had higher levels of proinflammatory cytokine IL1β (Figure 5D). This effect was more pronounced in infiltrating monocytes compared with microglia which is consistent with the relationship between autophagy deficiency and proinflammatory phenotype polarization after TBI.^16^

**Figure 5.**
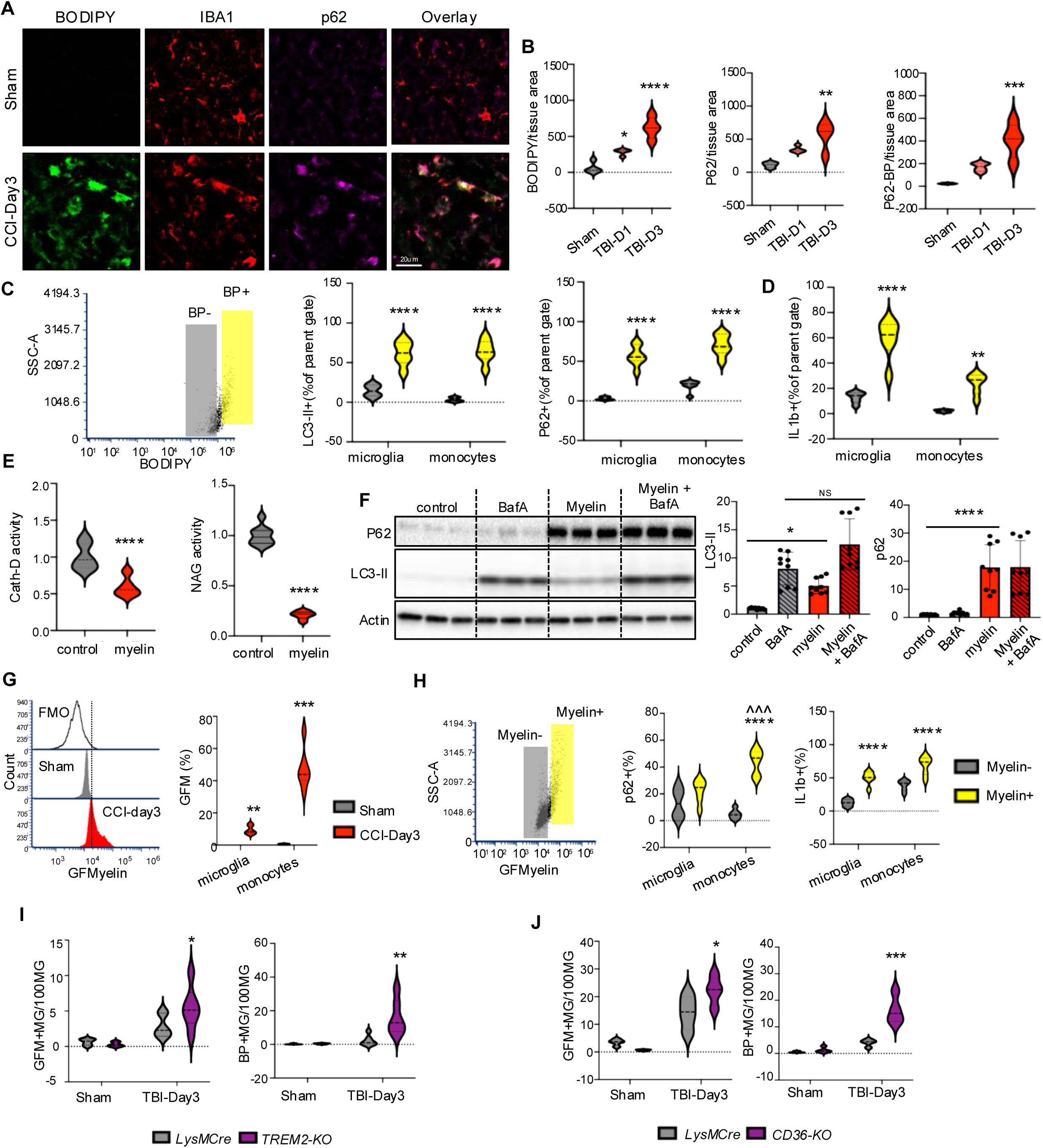
Myelin phagocytosis leads to inhibition of lysosomal function and autophagy after TBI. (A) Immunostaining demonstrating colocalization between lipid accumulation (BODIPY, green) and inhibition of autophagy (p62, magenta) in microglia/macrophages (IBA1, red) in cortical sections from TBI vs sham cortex (20X). (B) Quantification of data from panel A. n = 5 mice/group; * P < 0.05, ** P < 0.01, *** P < 0.001, **** P < 0.0001 (one-way ANOVA with Tukey’s). (C) Flow cytometry scatter plot representing gating strategy dividing TBI monocytes into BODIPY ^hi^ and BODIPY ^lo^ sub-populations. (D) Flow cytometry-based comparisons of autophagy (LC3 and p62, left and middle) and inflammatory (IL1β, right) markers in TBI microglia and monocytes with high (BODIPY^+^) vs low (BODIPY^-^) lipid accumulation. n = 6 mice/group; * P < 0.05. ** P < 0.001, **** P < 0.0001 (two-way ANOVA with Tukey’s). (E) Quantification of activity for lysosomal enzymes Cathepsin-D and NAGLU in BMDMs treated with purified mouse myelin (100 µg/ml – 24 hours) vs control. Values are mean ± SEM, n = 9 (3 replicates of 3 independent experiment); **** P < 0.0001 (Student’s t-test). (F) Western blots and quantification of autophagy markers p62 and LC3-II in BMDM treated with myelin vs control. Where indicated, Bafilomycin A (50 nM) was added for last 4 hours. Bar graphs are mean ± SEM, n = 9 (3 replicates of 3 independent experiments); * P < 0.05, **** P < 0.0001 (two-way ANOVA with Tukey’s). (G) Flow cytometry-based quantification of myelin (GreenFluoroMyelin, GFMyelin) accumulation in microglia and monocytes. Left - histogram demonstrating GFM signal shift from FMO in sham and TBI microglia. Right – quantification of GFM^+^ microglia and monocytes in TBI vs sham. n = 6 mice/group; * P < 0.05. ** P < 0.001, ***P<0.001 vs sham (Student’s t-test). (H) Flow cytometry scatter plot demonstrating identification of myelin^+^ and myelin^-^ monocyte sub-populations (left) and comparisons of autophagy marker p62 (center) and inflammation marker IL1β (right) in TBI microglia and monocytes with high vs low myelin accumulation. n = 6 mice/group. (I-J) Flow cytometry comparisons of myelin (GFMyelin, left plots) and neutral lipid (BODIPY, right plots) accumulation in mice deficient for *Trem2* (I) or conditionally deficient for *Cd36*. (J) vs corresponding WT controls. n = 6 mice/group. (two-way ANOVA with Tukey’s).

**Figure 6.**
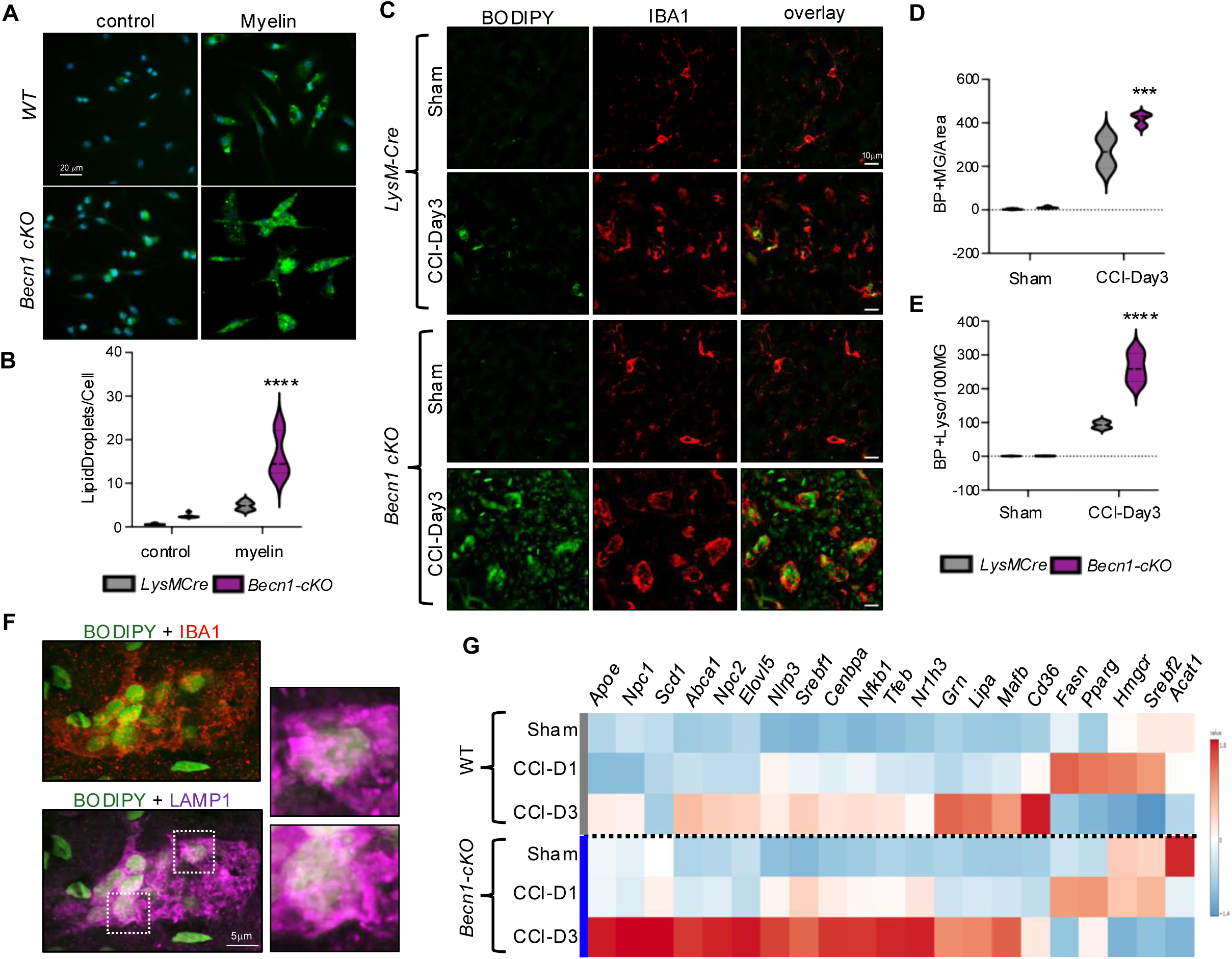
Autophagy deficiency exacerbates lipid accumulation and lipid metabolism reprograming. (A) BODIPY staining imaging of BMDMs (20x, scale bar 25 µm) from *LysMCre* (wild type) or *Beclin1-cKO* mice, following myelin treatment (100 µg/ml, 24 hours). (B) Quantification of data from panel A. n = 6 (3 replicates of 2 independent experiments); **** P < 0.0001 vs *LysMCre* (two-way ANOVA with Tukey’s). (C) Immunostaining demonstrating increased lipid (BODIPY, green) accumulation in microglia/macrophages (IBA1, red) in cortical sections of *Becn1-cKO* mice exposed to TBI as compared to *LysMCre* (wild type) mice (20X). (D-E) Quantification of IBA^+^ cell also positive for BODIPY per 100 IBA+ cells (20X images quantified in D) and colocalization of BODIPY and LAMP1 (60X images quantified in E). n = 4 mice/group; ** P < 0.01 vs *LysMCre* (two-way ANOVA with Tukey’s). (F) Confocal images demonstrating accumulation of neutral lipids (BODIPY, green) in lysosomes (LAMP1, magenta) of microglia (IBA1, red) in *Becn1-cKO* TBI brain sections. (G) RT-qPCR based heatmap comparing gene expression changes in lipid markers in peri-lesion cortical tissues from *LysMCre* (wild type) and *Beclin1-cKO* sham, TBI-day1, and TBI-day3 mice; n = 6 mice/group.

TBI causes generation of abundant myelin debris.^46^ We hypothesized that phagocytosis of fragmented myelin may contribute to lipid accumulating and lysosomal dysfunction after TBI. To confirm the effect of phagocytosed myelin on monocyte lysosomes, we isolated mouse bone marrow derived macrophages (BMDM)^47^ and exposed them to purified mouse myelin.^48^ We observed lysosomal accumulation of myelin, which was prevented by phagocytosis inhibitor Dynasore (Figure S5A-B). Additionally, we observed lipid droplet accumulation similar to that observed after TBI (Figure S5C). Consistent with the ability of phagocytosed myelin to inhibit lysosomal function, myelin treatment significantly decreased activity of lysosomal enzymes CathepsinD and N-acetyl-glucosaminidase in BMDM (Figure 6E). Furthermore, myelin led to inhibition of autophagy, as demonstrated by LC3-II flux assay and accumulation of p62 (Figure 5F, Figure S5D). We used flow cytometry to demonstrate accumulation of myelin (FluoroMyelin) in both resident microglia and infiltrating monocytes after TBI (Figure 5G). Similar to neutral lipid accumulation, myelin accumulation was more pronounced in infiltrating monocytes as compared to resident microglia. Furthermore, within TBI brain, microglial and monocyte cells with high levels of accumulated myelin (Myelin+) displayed higher extent of autophagy inhibition (p62) and higher inflammation (IL1β, Figure 5H). These data support the hypothesis that phagocytosis of myelin generated after TBI causes lysosomal lipid accumulation and inhibition of lysosomal function, which in turn leads to inhibition of autophagy.

Several receptors, including TREM2 and the scavenger receptor CD36, have been implicated in phagocytosis of myelin in different models.^23,49^ We used mice with global deficiency for *Trem2* (*Trem2^-/-^*) or conditional deficiency for *Cd36* in microglia and monocytes (*LysM-cre/Cd36-flox*) to determine their function in myelin phagocytosis and regulation of lipid accumulation after TBI. Surprisingly, both *Trem2* and *Cd36* deficient microglia and monocytes showed increase in myelin phagocytosis and neutral lipid accumulation after TBI (Figure 5I-J). However, the inflammatory responses for the two genotypes where not the same, with TNFα levels significantly higher in *Cd36* but not *Trem2* knock out microglia and monocytes as compared to wild type. IL1β levels were not altered in either genotype (Figure S6A-B).

### Autophagy deficiency exacerbates lipid accumulation and lipid metabolism reprogramming

Our data indicate that accumulation of myelin-derived lipids in the lysosomes leads to inhibition of autophagy after TBI. Autophagy has been also shown to participate in lipid homeostasis through lipophagy, which targets lipid droplets for lysosomal degradation and promotes cellular lipid efflux over storage.^45,50,51^ We hypothesized that autophagy deficiency would exacerbate lipid accumulation after TBI, creating a pathological feedback loop. We tested this hypothesis *in vitro* by exposing BMDMs from *LysM-cre/Becn1-flox* (*Becn1-cKO*) and *LysM-Cre* control mice to purified myelin and assessing lipid accumulation by BODIPY staining. We observed more lipid accumulation in *Becn1-cKO* BMDMs as compared to control both in the absence and presence of myelin, suggesting that autophagy deficiency is sufficient to promote lipid accumulation (Figure 6A-B). To test this *in vivo*, we subjected *Becn1-cKO* and *LysM-cre* control mice to CCI, followed by assessment of lipid accumulation at 3 days after injury. We observed significantly higher number of BODIPY^+^ microglia and macrophages (IBA1^+^) in brain tissue of injured *Becn1-cKO* as compared to *LysM-cre* control mice (Figure 6C-D). Staining for lipid droplet protein Perlipin 3 (PLIN3) demonstrated exacerbated lipid droplet formation in autophagy deficient monocytes after TBI (Figure S6C). *Becn1-cKO* mice also showed more lipid accumulation within lysosomes (Figure 6E-F). Finally, real-time qPCR analyses of perilesional cortex demonstrated exacerbated upregulation of lipid handling genes in injured *Becn1-cKO* as compared to *LysM-cre* mice, suggesting exacerbated reprograming of lipid metabolism after TBI (Figure 6G).

## Discussion

Recent data indicate an important contribution of altered lipid metabolism to microglial dysfunction during brain aging and in age-related neurodegenerative diseases such as AD.^1,5–7,10,52^ Much less is understood about the interactions between brain lipid environment, lipid metabolism and neuroinflammation during acute response to injury. Our data demonstrate that TBI leads to neutral lipid accumulation and rapid reprograming of lipid metabolism in microglia, brain associated macrophages and monocytes infiltrating the brain after injury-induced BBB disruption.^12^ Unlike aging and AD, where lipid reprograming is restricted to specific microglial sub-populations,^35^ after TBI virtually all microglia and monocytes in the ipsilateral hemisphere are affected to at least some extent. This likely reflects the overall breadth and intensity of microglial and monocyte inflammatory response after acute injury. These data support the notion that inflammatory and lipid metabolism responses are intimately linked, both under chronic and acute conditions.

While TBI-induced lipid metabolism reprograming was most pronounced in BAM and DAM populations, our data also revealed that homeostatic microglia responded more strongly as compared to surveillance microglia.^35^ The latter were defined as microglia enriched in the uninjured brain and expressing homeostatic as well as some activation markers. It is not clear why these populations may have attenuated lipid metabolic responses to injury as compared to homeostatic microglia. One possibility is that higher baseline activity of surveillance microglia makes them less prone to exaggerated responses to additional stimuli. Another, that homeostatic versus surveillance populations may be differentially distributed in respect to the injury site.^16^ Additional studies, such as spatial transcriptomic analyses of the injured brain will be necessary to differentiate between these possibilities.

In addition to demonstrating wide-spread lipid metabolism reprogramming after TBI, we used multi-omics approach to identify specific microglial (DAM_3) and monocyte (BAM_7) populations with exacerbated lipid metabolism changes and characterized by accumulation of neutral storage lipids such as cholesteryl esters and triglycerides. Inflammatory signaling through Toll-like receptor (TLR) activation has been previously shown to affect the lipidome of macrophages *in vitro*, with TLR4 stimulation specifically leading to neutral lipid accumulation and upregulation of triglyceride synthesis.^2^ This suggests that the lipidomic remodeling observed post-TBI may be partly a downstream consequence of TLR signaling triggered by damage-associated molecular patterns (DAMPs) released during brain injury. However, we also observed significant differences. Our analyses indicate that after TBI lipid handling pathways are affected to a greater extent as compared to lipid synthesis. This was apparent both in overall analyses of microglial/monocyte responses as well as in lipid accumulating populations such as BAM_7 and DAM_3. This suggests that defects in lipid handling and efflux rather than activation of lipid synthesis may be the predominant mechanism leading to microglial/monocyte lipid accumulation after TBI. The reasons for this differential reprograming are not clear. While it is possible that it could reflect differences between cell types, overall reprograming patterns in monocytes infiltrating the brain after TBI resembled those observed in microglia. Additionally, while TLR activation by DAMPs plays an important role in activation of interferon signaling pathways,^53,54^ our analyses indicate that microglial/monocyte populations accumulating lipid after TBI are distinct from those with most pronounced activation of interferon responses. An alternative possibility is that the lipid-rich environment of the brain parenchyma interacts with pro-inflammatory stimuli such as TLR ligands to modulate cellular responses. This is supported by our and other data demonstrating that myelin phagocytosis leads to cellular lipid accumulation and altered expression of lipid metabolism and inflammatory genes in monocytes and microglia.^25,55^ While further analyses will be necessary to decipher these interactions, together, our findings and those of Hsieh et al.^2^ point to lipidome remodeling as a conserved response mechanism in macrophage-lineage cells across different inflammatory contexts, reinforcing the role of lipid metabolism as both a marker and mediator of inflammation.

Our data indicate that after TBI monocytes show more pronounced lipid accumulation and lipid metabolism reprograming as compared to activated microglia. Infiltrating monocyte-derived macrophages show stronger phagocytic uptake of apoptotic neurons and myelin fragments, which are rich in phospholipids, sphingolipids and cholesterol. Once internalized, these lipids are processed in the lysosomes and eventually stored in lipid droplets, especially when catabolic or efflux pathways are overwhelmed or dysfunctional. Additional factor contributing to the more pronounced phenotype in the infiltrating cells may be that unlike microglia, they have not been exposed to the unique brain lipid milieu. Recent studies show that infiltrating macrophages in several models of CNS injury can exhibit a “foamy” phenotype marked by lipid accumulation, increased levels of lipid droplet-associated proteins like PLIN2 and SOAT1, and dysregulation of cholesterol efflux.^25,56,57^ The impaired lipid handling may also be further amplified by environmental cues in the TBI lesion such as oxidized lipids, which potentiate scavenger receptor activity and lipid droplet formation.^58^ Intriguingly, HIF-1 signaling was one of the top hits in DAM_3 pathway analysis, which is in agreement with a recent observation that lipoprotein-derived fatty acids generated through LIPA activity stabilize HIF-1α independent of hypoxia.^59^ Therefore, the enhanced lipid burden in infiltrating macrophages may reflect both higher phagocytic uptake and lower capacity for intracellular lipid metabolism in these recruited cells.

CD36 and TREM2 receptors have been both implicated in myelin phagocytosis^23,56^. The unexpected increase in both myelin and lipid accumulation observed in *Cd36* and *Trem2* deficient microglia and monocytes following brain trauma may reflect redundancy and compensatory mechanisms among scavenger receptors. Additionally, while both CD36 and TREM2 are recognized for their roles in lipid uptake, emerging evidence suggests they also participate in regulation of cellular lipid processing and efflux.^23^ *Trem2*-deficient microglia have been reported to exhibit impaired clearance of cholesterol, which was associated with downregulation of genes involved in lipid catabolism and efflux, such as *Apoe* and *Abca1*, leading to accumulation of cholesteryl esters and lipid droplets.^60^ Similarly, inhibition of CD36 has been shown to reduce activity of LXR and PPAR and promote inflammation in response to myelin treatment *in vitro*^23^. This is consistent with our data showing exacerbation of some inflammatory responses in *Cd36* but not *Trem2* deficient mice. We expect that in the absence of CD36 or TREM2, other receptors may partially compensate for lipid uptake, but not for the downstream regulation of lipid processing and efflux pathways.

Our imaging and lysosome-specific lipidomic data indicate that after TBI neutral lipids including cholesterol esters and triglycerides accumulate in both, lipid droplets and lysosomes. Additionally, we observed increase in lysosomal ceramides and sphingomyelins. Cholesterol, ceramides and sphingomyelins are all major components of myelin, and their lysosomal accumulation supports myelin debris phagocytosis as a source of lysosomal lipids after TBI. Triglycerides, however, are not enriched in myelin. Our proteomic analyses revealed apolipoprotein E (APOE) as the top protein accumulating in lysosomes after TBI. As APOE is the major neutral lipid carrier in the brain,^61,62^ this suggests that triglycerides and potentially some cholesterols and may be derived from lipoprotein phagocytosis. Another protein enriched in TBI lysosomes was ANXA5, which mediates phagocytic clearance of apoptotic cells and debris through direct interaction with phosphatidylserine on their membranes.^63^ In addition to its role as lipid carrier, APOE is well-established in modulation of lipid metabolism through the activation of LXR.^64^ Consistently, our scRNA-seq identified the LXR (NR1H3) pathway as the most significant regulatory network in the lipid-accumulating BAM_7 cluster. This convergence of multi-omics findings highlights the LXR pathway as a central regulator of lipid handling and inflammatory responses in monocytes following injury and suggests it as a compelling therapeutic target in TBI.

In peripheral phagocytes including macrophages uptake of certain lipids including cholesterol can directly cause lysosomal dysfunction. The proposed mechanisms include inhibition of vesicle fusion due to altered membrane composition, and physical disruption of lysosomal membranes.^65,66^ This has been documented in tissues high in cholesterol such as atherosclerotic plaques and demyelinated lesions in multiple sclerosis, where accumulation of lipid droplets in foam macrophages and lipid-loaded microglia is associated with lysosomal dysfunction, defective phagocytosis, and inflammation.^48^ Our data indicate that, similarly, phagocytosis of myelin debris after TBI affects lysosomal function and leads to inhibition of autophagy in affected microglia and monocytes. We recently demonstrated that TBI causes inhibition of microglial and monocyte autophagy and that this inhibition is an important contributing factor to excessive and prolonged neuroinflammatory responses.^16,18^ However, the mechanisms leading to autophagy inhibition after TBI remained unknown. The data presented here indicate that lipid and especially myelin phagocytosis induces lysosomal dysfunction and, consequently inhibition of autophagy in the injured brain. In addition to its role in maintenance of protein and organelle homeostasis and regulation of inflammation, recent data indicate that autophagy also regulates lipid metabolism.^19,21,22^ Autophagy has been shown to specifically target and degrade lipid droplets through lipophagy, thus promoting lipid efflux over storage. Consistently, we observed more pronounced lipid metabolism reprograming and exacerbated cellular lipid accumulation in mice with microglia/monocyte specific autophagy defects. Our data suggest a pathological feedback loop, where lipid phagocytosis causes inhibition of autophagy-lysosomal function after TBI, which in turn exacerbates cellular lipid retention, reprograming and inflammation.

Our study highlights lipid metabolism imbalance within mononuclear phagocyte populations as a central hallmark of the innate immune response in the context of TBI. This dysregulation manifests as lipid droplet accumulation, altered expression of genes involved in lipid processing, and a shift toward pro-inflammatory metabolic programs. Importantly, these changes are not restricted to TBI but appear to be a convergent feature across diverse forms of acute CNS inflammation. Similar lipid accumulation in microglia and/or macrophages has been documented in neuroinflammatory contexts such as spinal cord injury,^57^ ischemic stroke,^56^ and multiple sclerosis,^25^ suggesting a shared mechanism by which innate immune cells present in the CNS adapt their lipid metabolism in response to inflammation. These findings underscore the potential of targeting lipid metabolism pathways as a unifying therapeutic approach across multiple acute CNS pathologies where innate immune dysfunction is a key contributor. Additionally, lipid droplet associated microglia (LDAM) are observed and play a pathological role during brain aging and in neurodegenerative diseases.^1,9^ History of CNS injury including TBI is a strong predisposing factor to AD and other AD-related dementias later in life.^67–69^ Since the lipid accumulating microglial and monocyte populations we identified in the TBI brain strongly resemble LDAM, we expect that they may contribute to the accelerated development of dementia after injury.

## Acknowledgements

This work was supported by R01NS115876 and R56AG081262 to MML. Confocal imaging was performed at the University of Maryland Center for Innovative Biomedical Resources Confocal Microscopy Core Facility, supported by S10OD026698. Flow cytometry was performed at the Flow Cytometry Shared Service at the University of Maryland Marlene and Stewart Greenebaum Comprehensive Cancer Center, supported by Maryland Department of Health’s Cigarette Restitution Fund Program (CH-649-CRF) and P30CA134274. RNA sequencing was performed by Maryland Genomics at the University of Maryland Institute for Genome Sciences.

## Author Contributions

Conceptualization: A.A.M.T., and M.M.L. Experiments: A.A.M.T., N.H., B.R.H., C.S., S.T., D.PH.N., S.A.K., Y.M., M.M.W., L.M., W.T.A., M.C.G., O.P.R., S.B., L.K.G., and J.T.G. Data analysis: A.A.M.T., B.R.H., M.M.W., W.T.A., J.W.J., and M.M.L. Writing-original draft: A.A.M.T. and M.M.L. Supervision: M.M.L., M.A.K., J.W.J., and S.A.A. Funding acquisition: M.M.L., M.A.K. and J.W.J.

---

**Figure S1.**
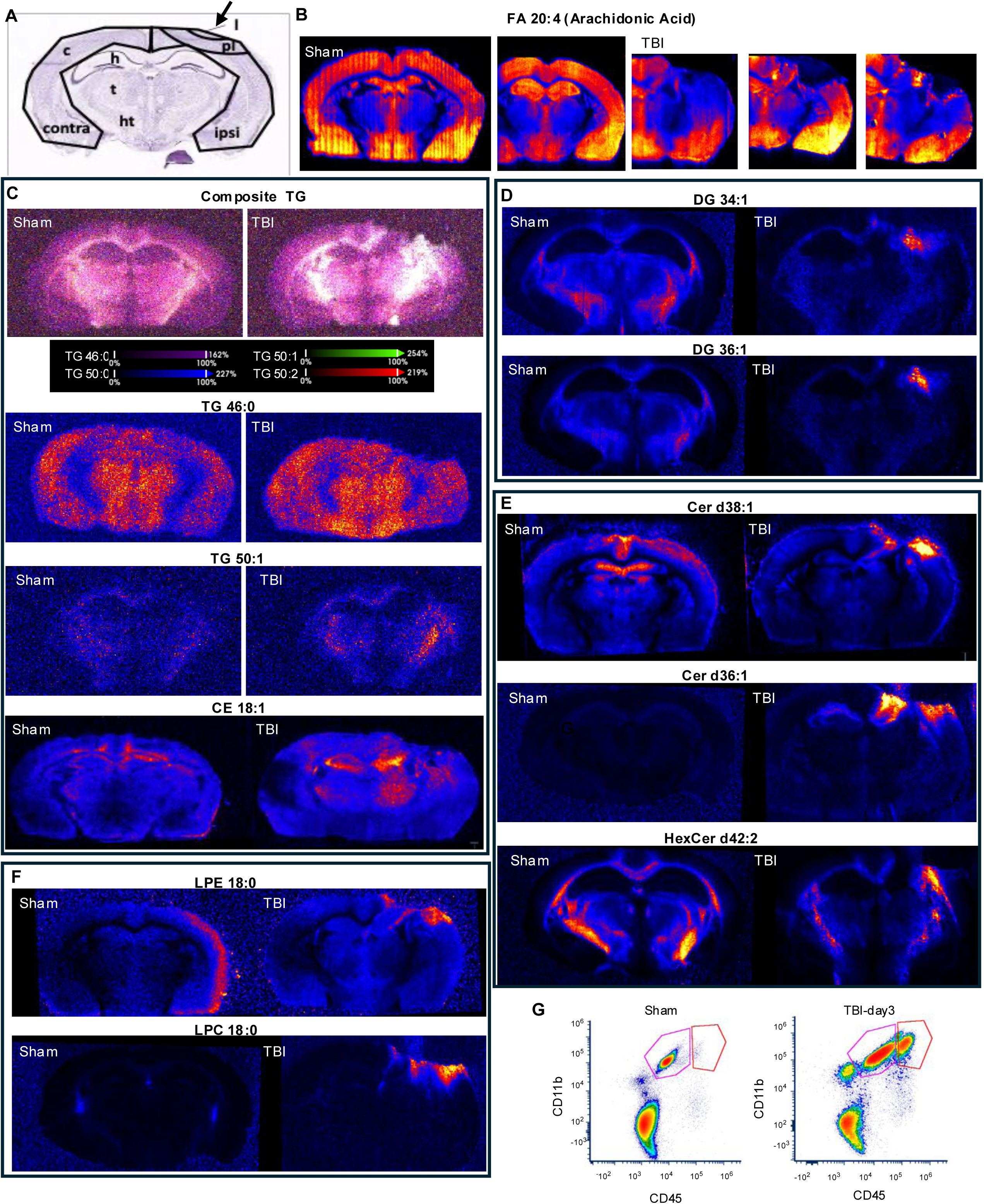
TBI leads to accumulation of neutral lipids in microglia and infiltrating macrophages. Related to Figure 1. (A) Schematic representation of a coronal mouse brain section illustrating spatial terminology used in this study. The ipsilateral hemisphere (ipsi) refers to the side of the brain directly impacted in the controlled cortical impact (CCI) model of TBI (lesion site (l) is indicated with arrow), while the contralateral hemisphere (contra) is the uninjured side. The perilesional (pl) cortex (c) denotes the cortical region immediately adjacent to the lesion core (above hippocampus (h)) representing tissue at risk for secondary injury and a key area of immune and metabolic response. This schematic serves as a reference for the anatomical orientation of tissue punches, imaging regions, and data interpretation throughout the manuscript. Thalamus (t), hypothalamus (ht). (B-F) Desorption Electrospray Ionization Mass Spectrometry Imaging (DESI-MSI) of coronal mouse brain slices, sham vs TBI day 3 showing spatial distribution of different lipid species. (B) Arachidonic acid (FA 20:4) in two sham (left) and three TBI (right) animals demonstrating reproducibility. (C) Neutral lipids including composite of four triglyceride (TG) species (top), two of the component TG (TG 46:0 and TG 50:1) and cholesteryl ester (CE 16:0). (D) Diacyl glycerols (DG 34:1 and DG 36:1). (E) Ceramides (Cer d38:1 and Cer d18:1/18:0) and hexosylceramide (HexCer d42:2). (F) Lyso-phospholipids including lysophosphatidylethanolamine (LPE 18:0) and lysophosphatidylcholine (LPC 18:0). (G) Flow cytometry scatterplot demonstrating gating strategy for identifying microglia (CD11B+CD45int) and monocytes (CD11B+CD45hi) in sham (left) and TBI (right).

**Figure S2.**
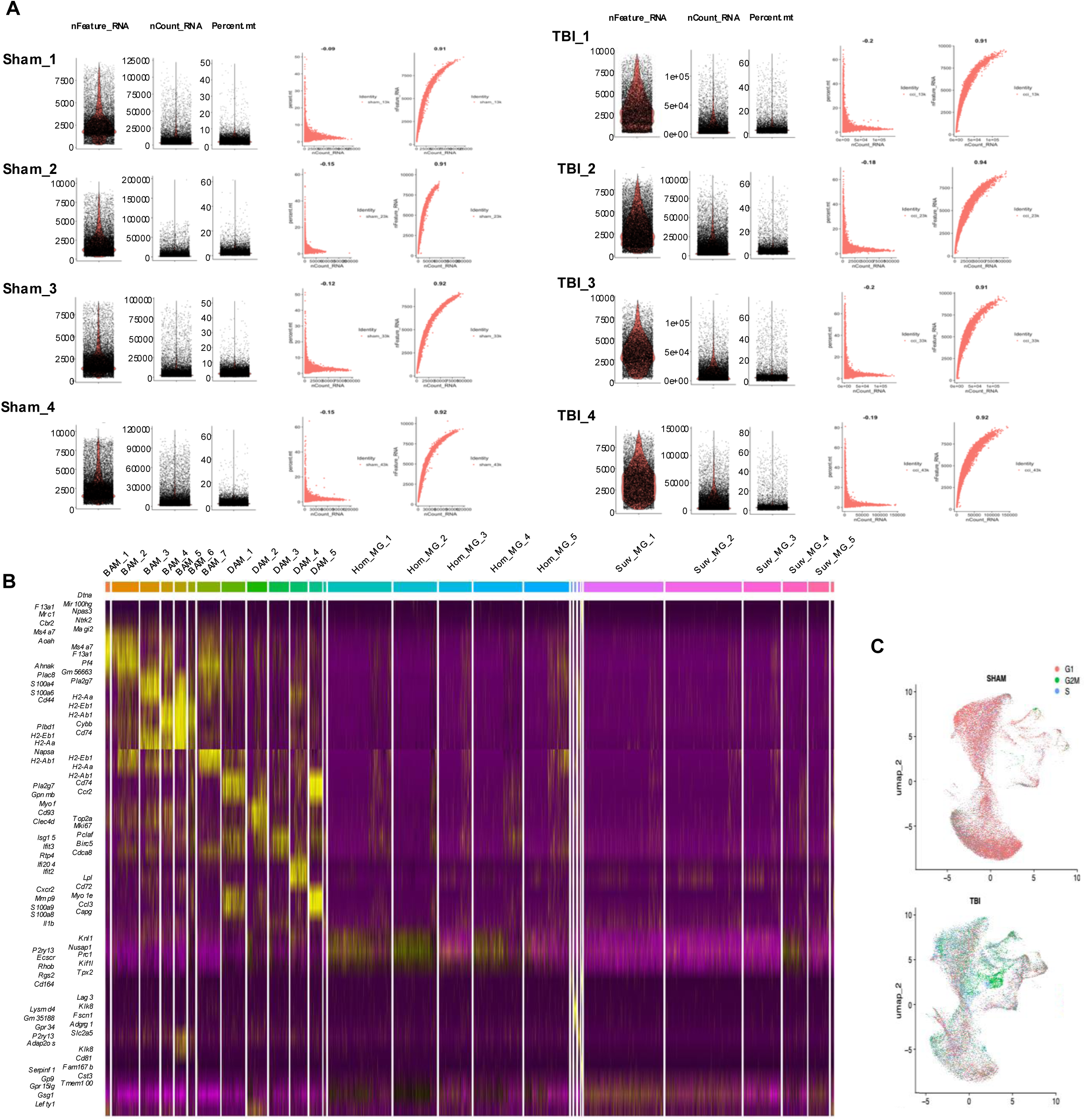
TBI causes pronounced lipid metabolism reprogramming in microglia and macrophages. Related to Figure 2. (A) Quality control plots for each of sham 1-4 (left) and TBI 1-4 (right) samples used for scRNA-seq analyses. Left – three violin plots visualizing nFeature_RNA, nCount_RNA, and percent.mt QC metrics. Right – two FeatureScatter() generated plots of nCount_RNA vs perent.mt and ncount_RNA vs nFeature_RNA showing overall complexity and mitochondrial stress across cells and samples. Together, these panels confirm the technical consistency and biological comparability across the four individual samples in both the sham and TBI groups. All samples underwent identical QC filtering thresholds, and no individual replicate displaying outlier behavior in these metrics. (B) Heatmap displaying top 5 differentially expressed genes for each cluster, selected based on highest average log fold-change using Seurat’s FindAllMarkers() function. Gene expression values were scaled across cells and centered to highlight relative expression patterns. Each column represents a single cell, grouped by cluster identity, and each row corresponds to a marker gene. This visualization highlights cluster-specific transcriptional signatures, enabling annotation of distinct cell types and states. Marker genes were selected using a Wilcoxon rank-sum test with an adjusted p-value < 0.05 and minimum expression threshold across cells. Color scale indicates z-scored expression levels. (C) Cell cycle state classification of single-cell transcriptomes from sham and TBI cortex. Uniform Manifold Approximation and Projection (UMAP) plots showing cells color-coded by predicted cell cycle phase (G1, S, or G2/M) using Seurat’s built-in cell cycle scoring pipeline. Cell cycle phases were inferred based on canonical marker gene expression profiles. The analysis reveals a higher proportion of TBI-associated cells in S and G2/M phases, consistent with overall increased proliferative activity. This shift explains the observed elevation in nFeature_RNA values after TBI in panel A, and supports biological findings from downstream pathway enrichment analysis, which identified cell division-related terms as the most significantly upregulated in the TBI condition (Figure 2E).

**Figure S3.**
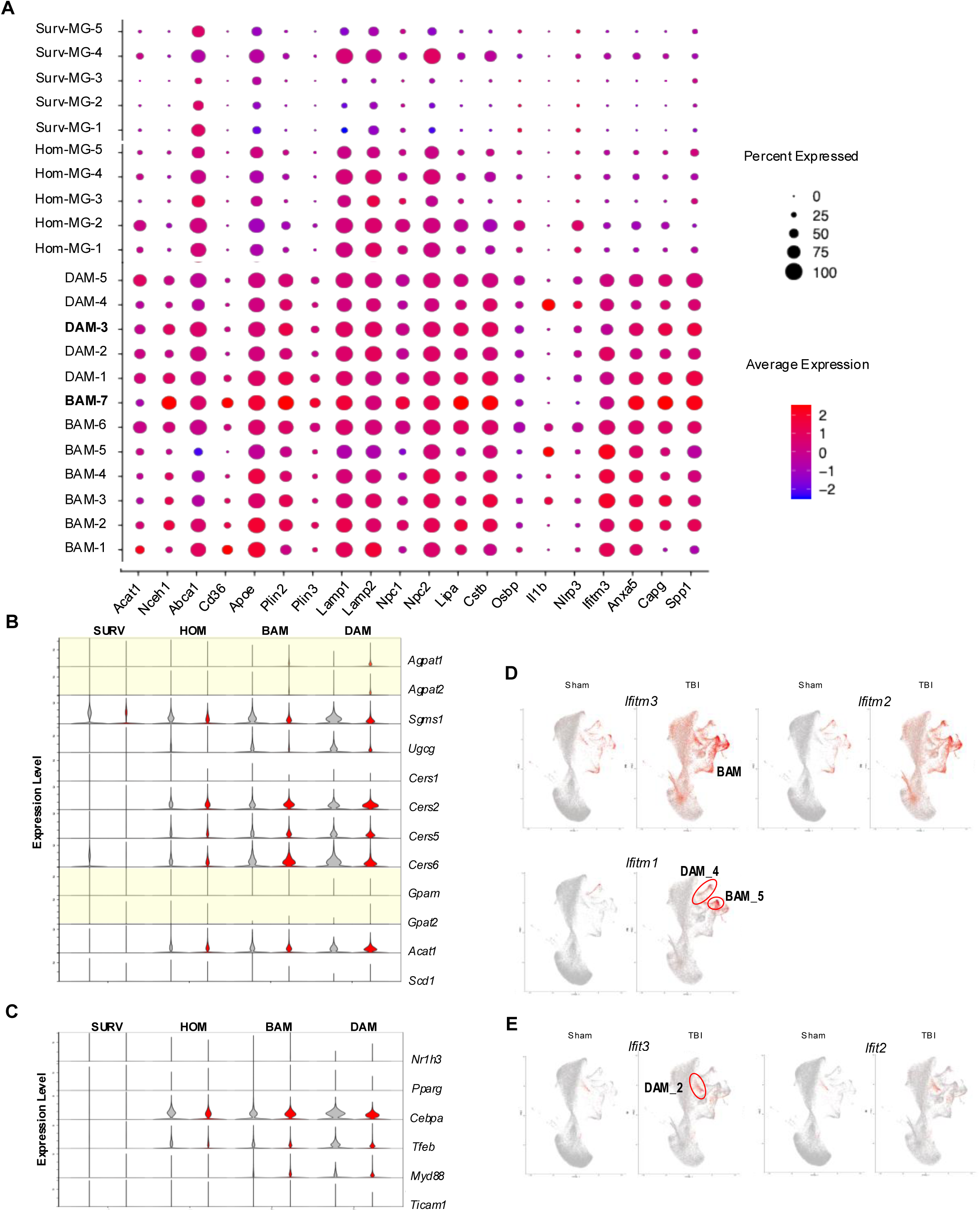
Lipid reprogramming is more pronounced in lipid-accumulating microglia and macrophages. Related to Figures 2 and 3. (A) Dot plot showing expression of selected lipid metabolism, lysosomal function and phagocytosis marker genes across major microglial and macrophage subtypes. Each dot represents the expression of a given gene (x-axis) in each cluster (y-axis) of all four main categories: homeostatic microglia (HOM), surveillance microglia (SURV), disease-associated microglia (DAM), and border-associated macrophages (BAM). Dot size indicates the percentage of cells in each cluster expressing specified gene; color intensity reflects the average scaled expression level (z-score) among expressing cells. (B) Violin plots (generated by ggplot() in Seurat) showing expression levels of lipid synthesis genes involved in the synthesis of triglycerides (highlighted in yellow: *Agpat1, Agpat2, Gpam*, and *Gpat*), sphingomyelin (*Sgms1*), ceramides (*Cers1*, 2, 5, and 6) and cholesterol (*Acat1* and *Scd1*). (C) Violin plots (generated by ggplot() in Seurat) showing expression levels of select transcription factors involved in lipid metabolism and TLR signaling. (D-E) Feature plot visualization of interferon-stimulated gene expression across all microglial and monocyte populations emphasizing the heterogeneous and cluster-specific activation of interferon-responsive genes following TBI. (D) *Ifitm3* and *Ifitm2* show elevated expression in both BAMs and DAMs, with a stronger signal in BAM subclusters. *Ifitm1* is primarily expressed in *DAM4* and *BAM5*, highlighting a subpopulation-specific interferon response. (E) *Ifit3* and *Ifit2* exhibit highly selective expression in *DAM2*.

**Figure S4.**
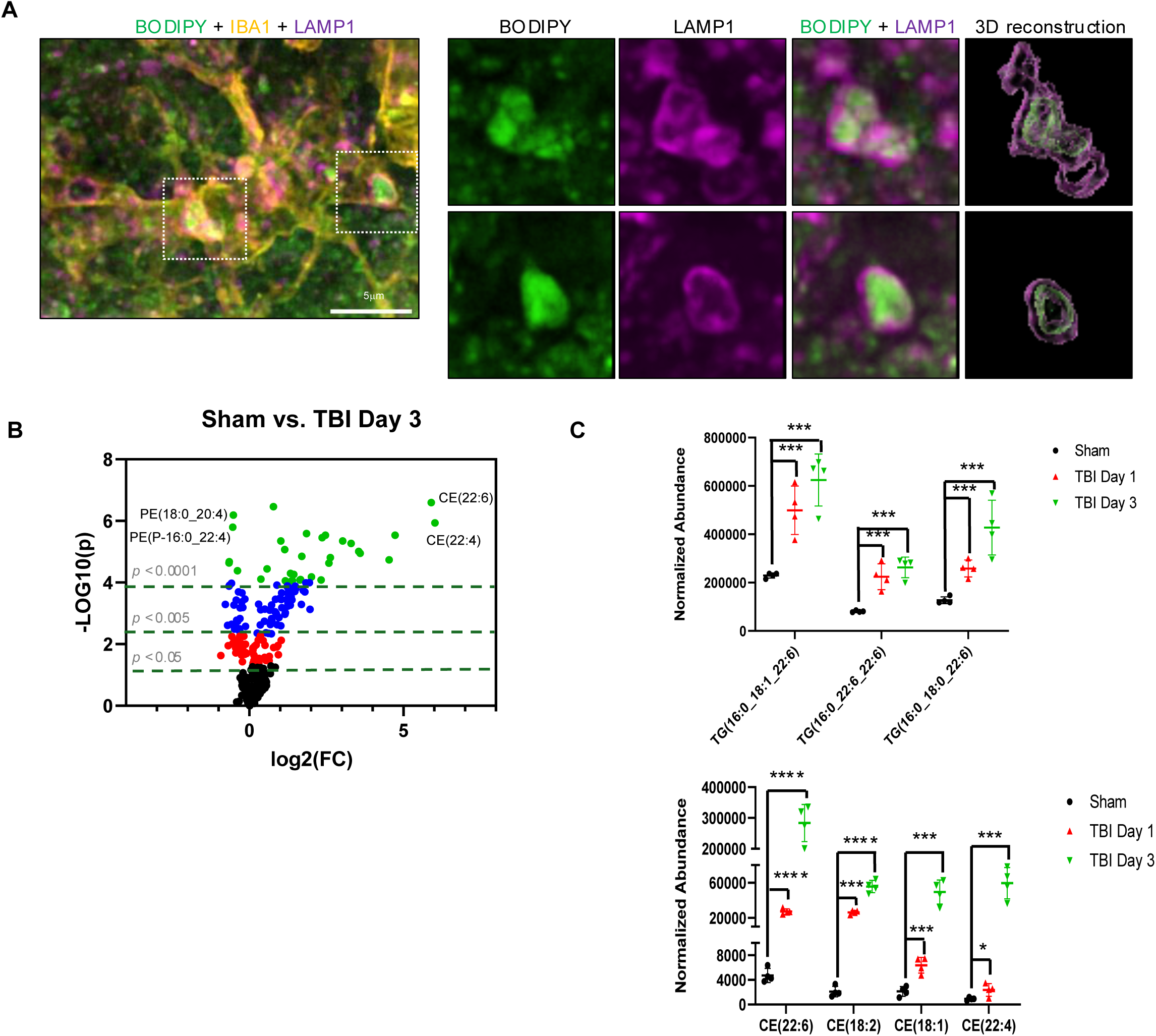
Lipid accumulates in the lysosomes of microglia and macrophages after TBI. Related to Figure 4. (A) Close up confocal image of active microglia after TBI and insets of single channels used for 3D reconstruction. (B) MetaboAnalyst generated volcano plot highlighting lipids with P < 0.05 (red), P < 0.005 (blue), and P < 0.0001 (green, based on t-test) in TBI (day 3) vs sham. Identity of selected lipid species is indicated. n = 4 mice/group. (C) Normalized abundance of selected differentially abundant triglyceride (TG, top) and cholesteryl ester (CE, bottom) species across experimental groups. Each point represents an individual sample; bars indicate mean ± standard deviation. Lipid species shown on the x-axis are among those with the highest fold-change and statistical significance in TBI vs Sham groups. * p <0.05, ** p < 0.01, *** p < 0.005, **** p < 0.0001 (one-way ANOVA with FDR correction).

**Figure S5.**
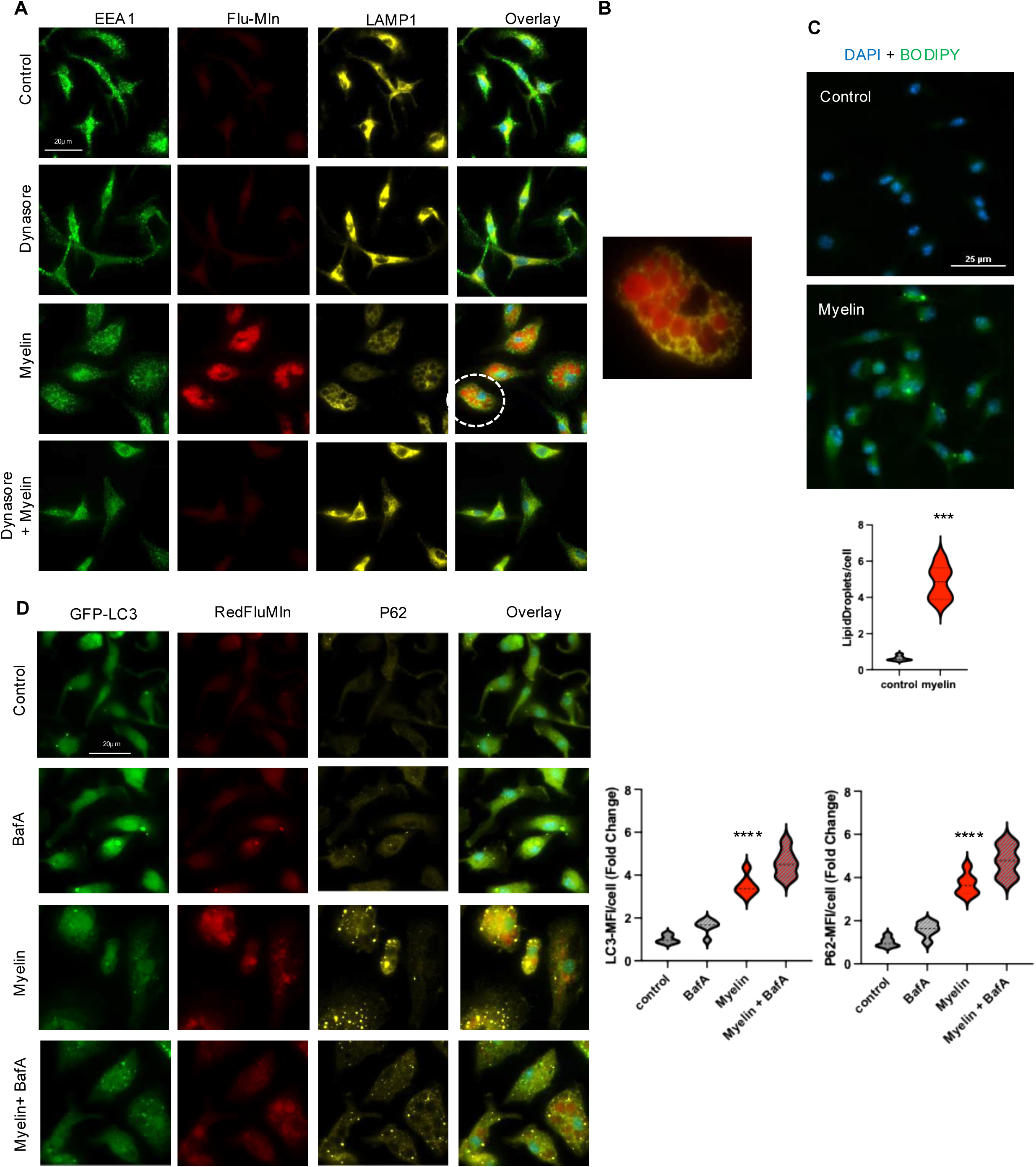
Myelin phagocytosis inhibits lysosomal function and autophagy after TBI. Related to Figure 5. (A) Immunofluorescence images demonstrating phagocytic myelin uptake and its subcellular localization to lysosomes in wild-type bone marrow–derived macrophages (BMDMs). BMDM were treated with vehicle (control), purified mouse myelin (100 μg/mL, 4 hr), Dynasore (10 μM, 4.5 hr), or Dynasore pre-treatment (30 min) followed by myelin (4 hr). Cells were stained for early endosomes (EEA1, green), phagocytosed myelin (RedFluoroMyelin, red), and lysosomes (LAMP1, yellow). BMDMs exhibited robust myelin uptake, which was prevented by Dynasore, confirming that myelin internalization is dynamin-dependent. (B) Inset from indicated are in panel A, demonstrating accumulation of myelin (RedFluoroMyelin) within LAMP1 lysosomes. (C) Images and quantification demonstrating that myelin treatment induced neutral lipid accumulation in wild-type BMDMs. BMDM were treated with vehicle (control) or myelin (100 μg/mL, 24 hr). P < 0.05 (unpaired *t*-test). (D) Immunofluorescence images and quantification demonstrating inhibition of autophagy flux in GFP-LC3–expressing BMDMs treated with myelin. GFP-LC3 BMDM were treated with vehicle or myelin (100 μg/mL, 24 hr), ± Bafilomycin A1 (50 nM for the final 4 hr). ****P < 0.001 vs control; values are mean ± SEM, n = 6 (3 replicates of 2 independent experiments); **** P < 0.0001 (two-way ANOVA with Tukey’s).

**Figure S6.**
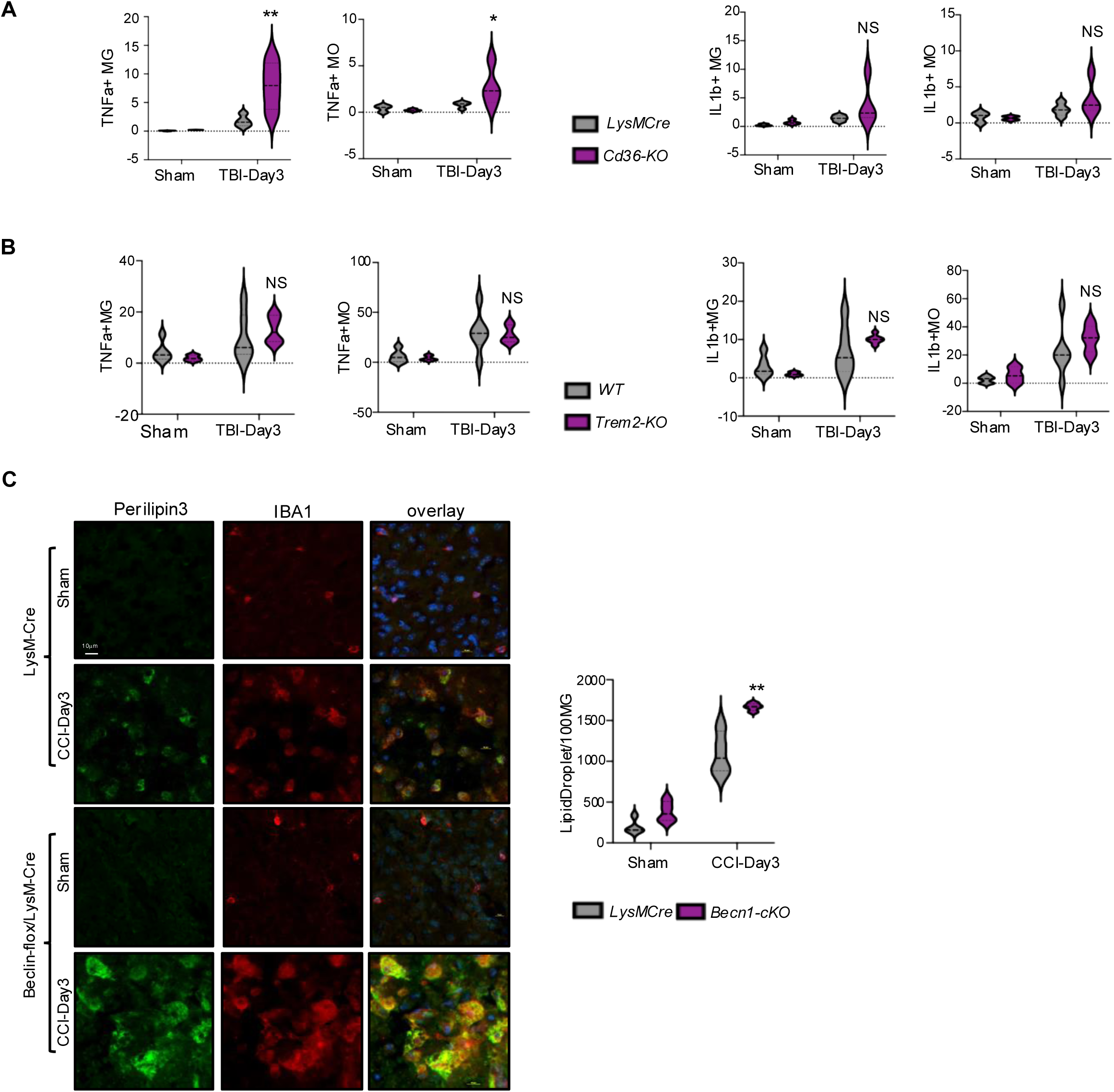
Autophagy deficiency exacerbates lipid accumulation and lipid metabolism reprograming. Related to Figures 5 and 6. (A-B) Flow cytometry quantification of inflammatory cytokine TNFa (left) and IL1b (right) expression in microglia and monocytes from *Cd36-* (A) and *Trem2*-(B) deficient as compared to corresponding WT control mice following TBI. n = 6 mice/group; *P < 0.05, **P < 0.01 vs corresponding WT (two-way ANOVA with Tukey’s). (C) Immunofluorescence images and quantification of demonstrating increased lipid droplet (Perilipin 3) formation in microglia/monocytes (IBA1) in *Beclin1-cKO* vs *LysMCre (WT)* mouse brains at day 3 post-TBI. n = 4 mice/group; **P < 0.01 vs *LysMCre* (WT) (two-way ANOVA with Tukey’s).

## STAR*METHODS

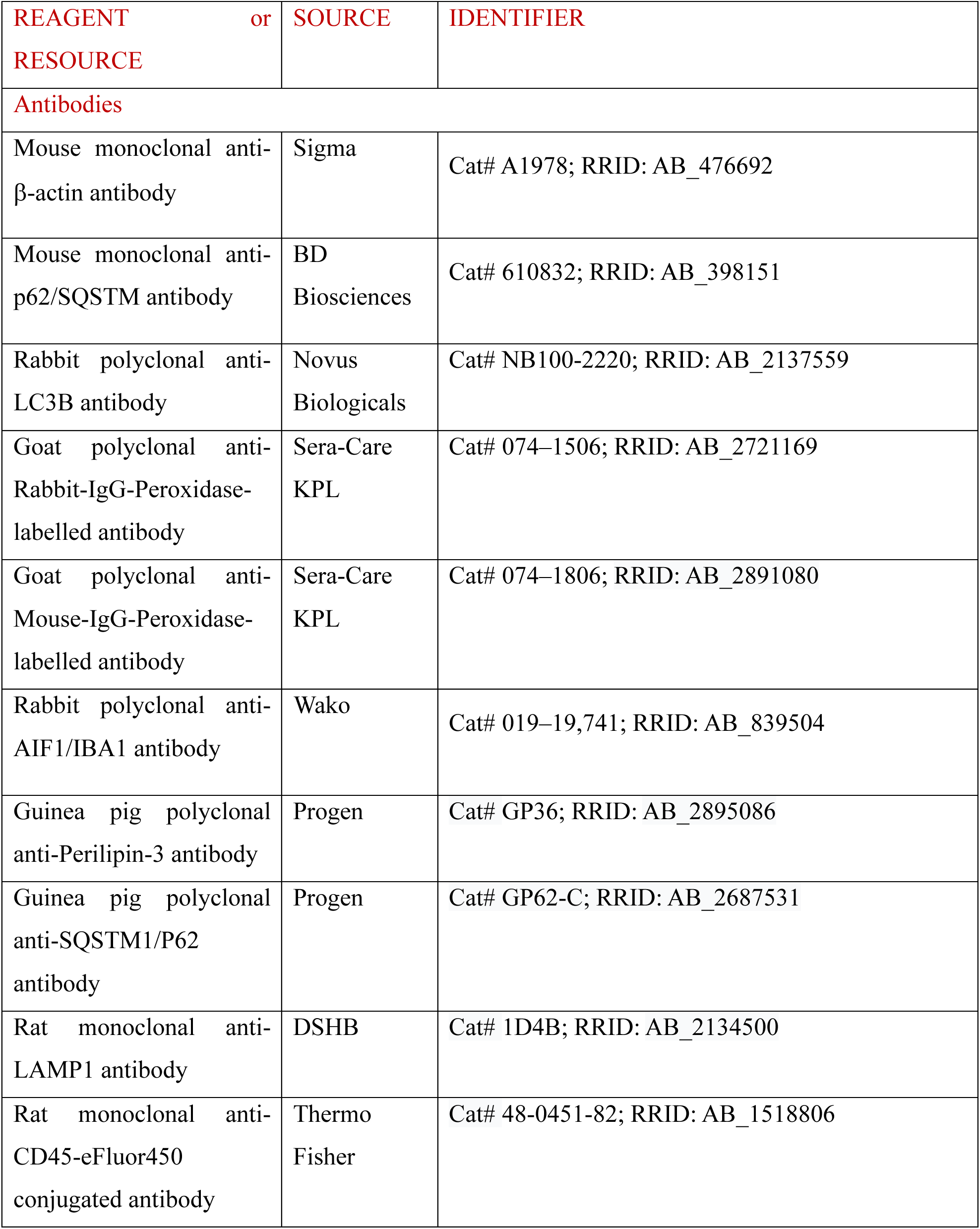
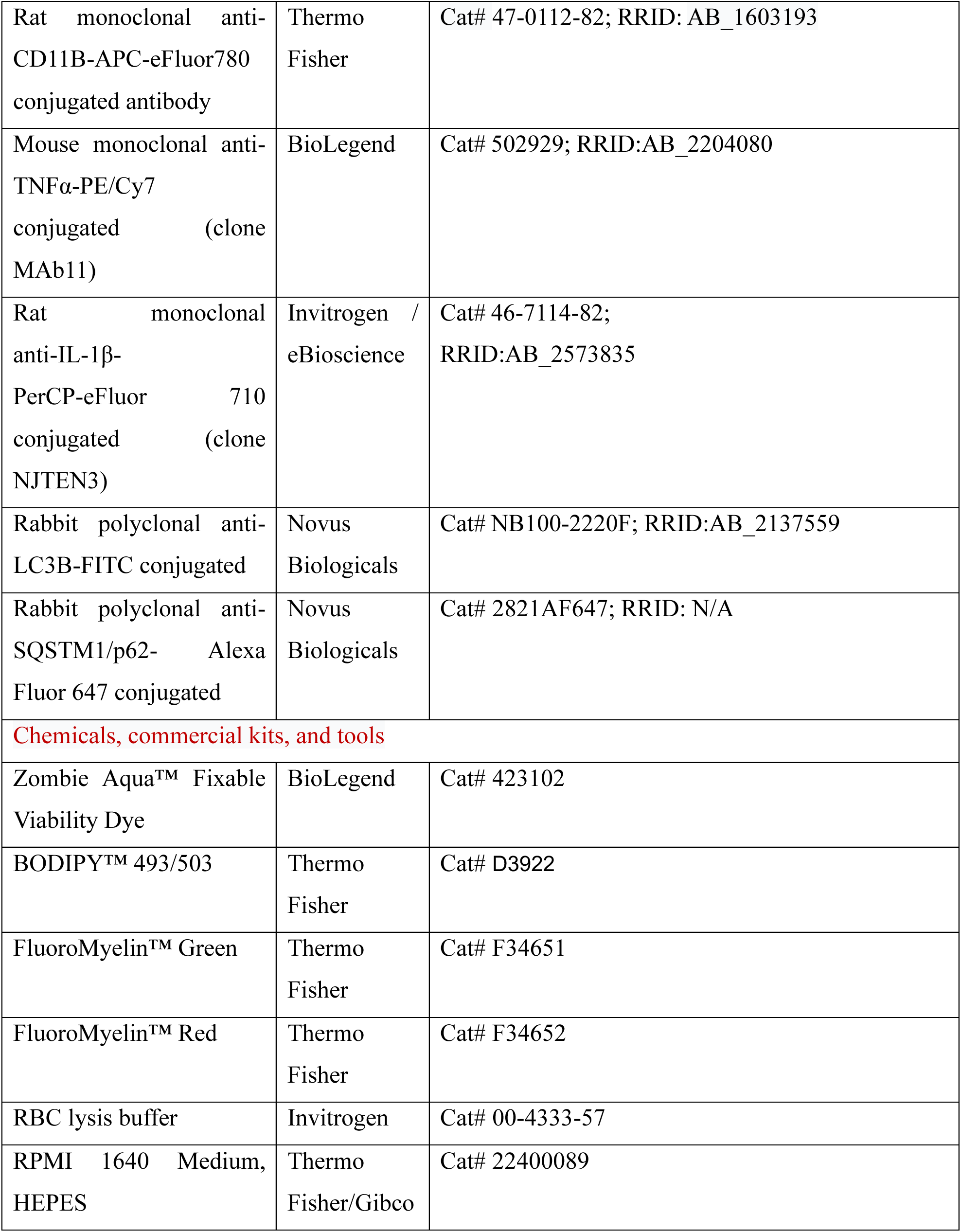
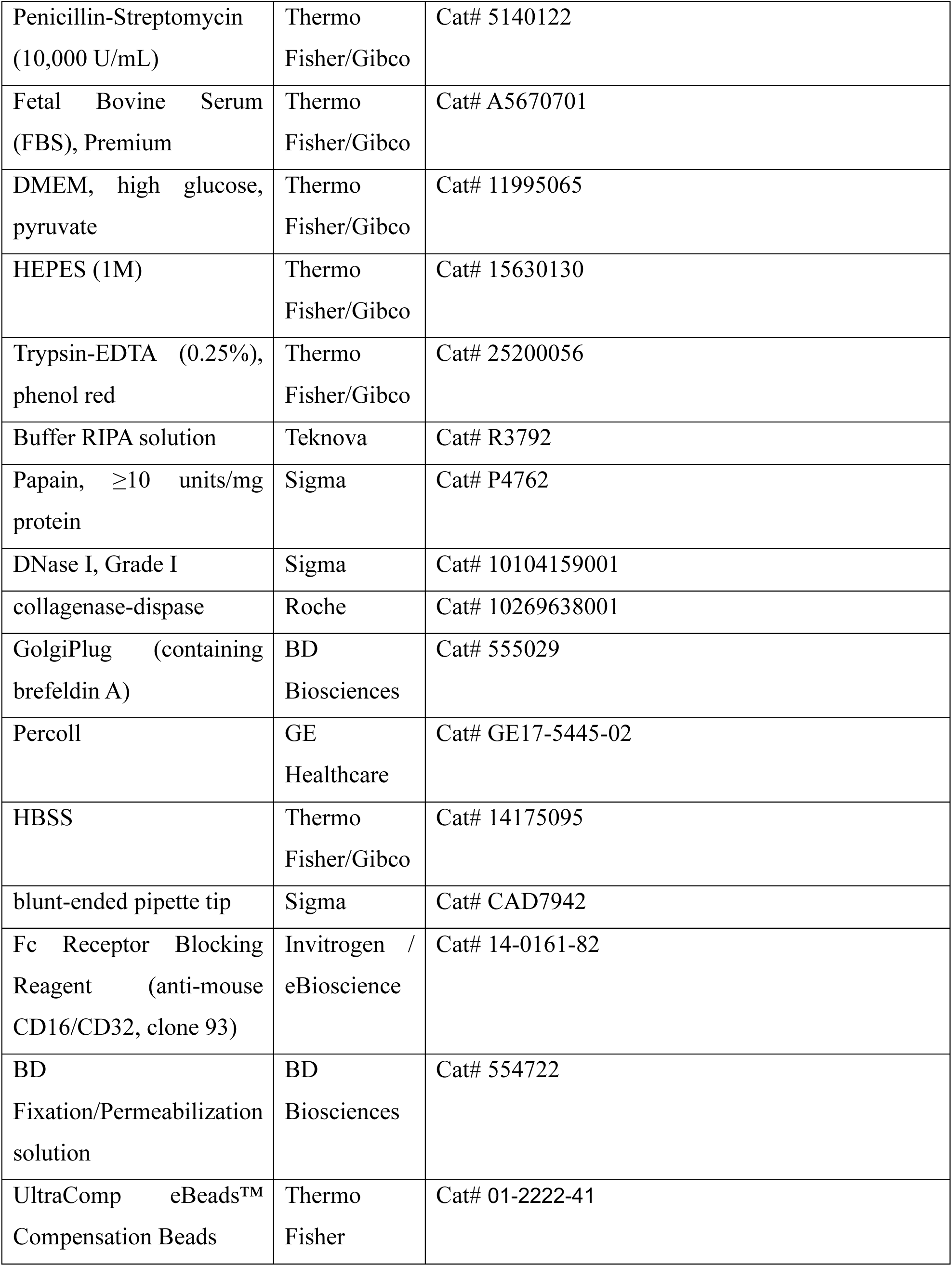
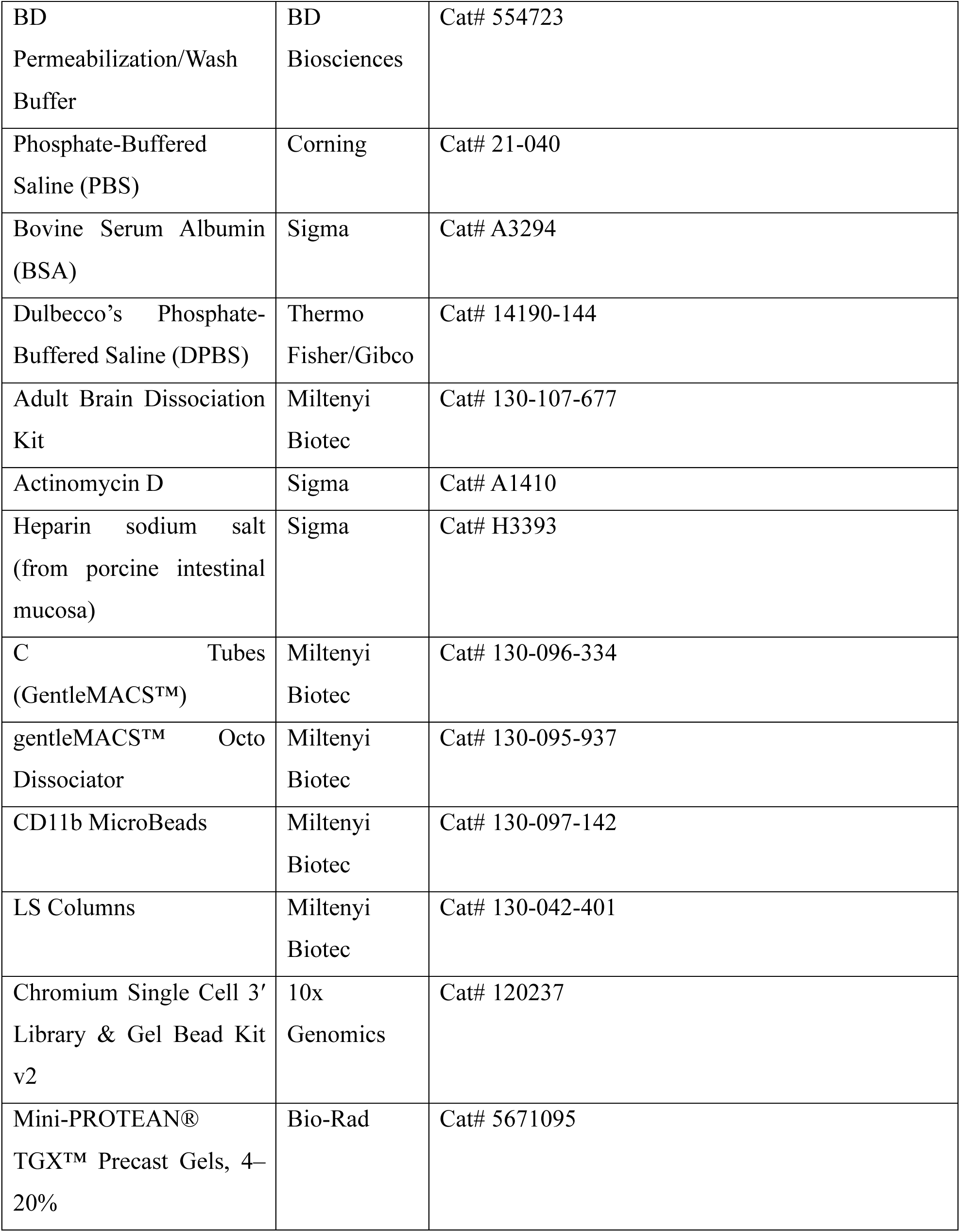
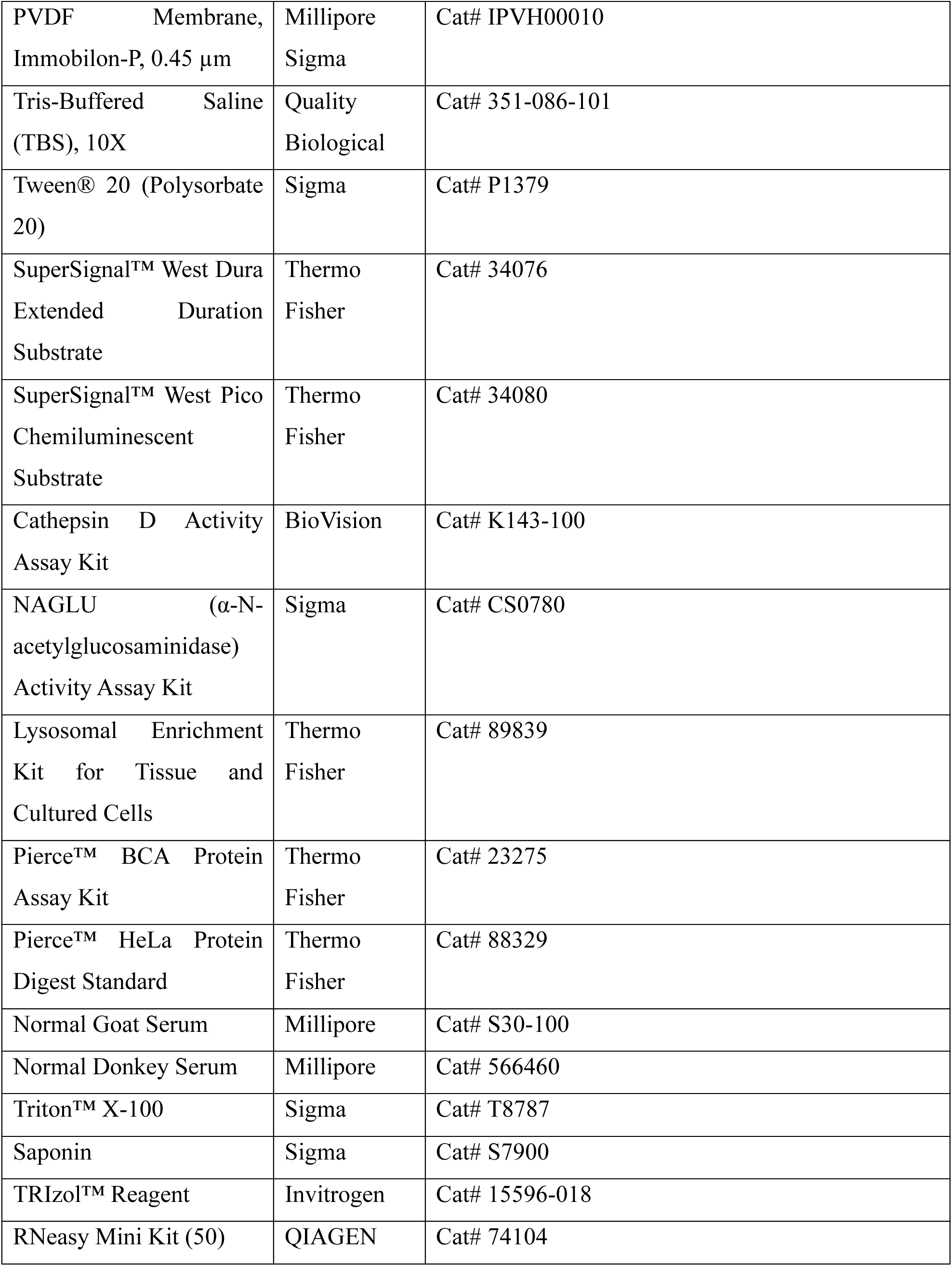
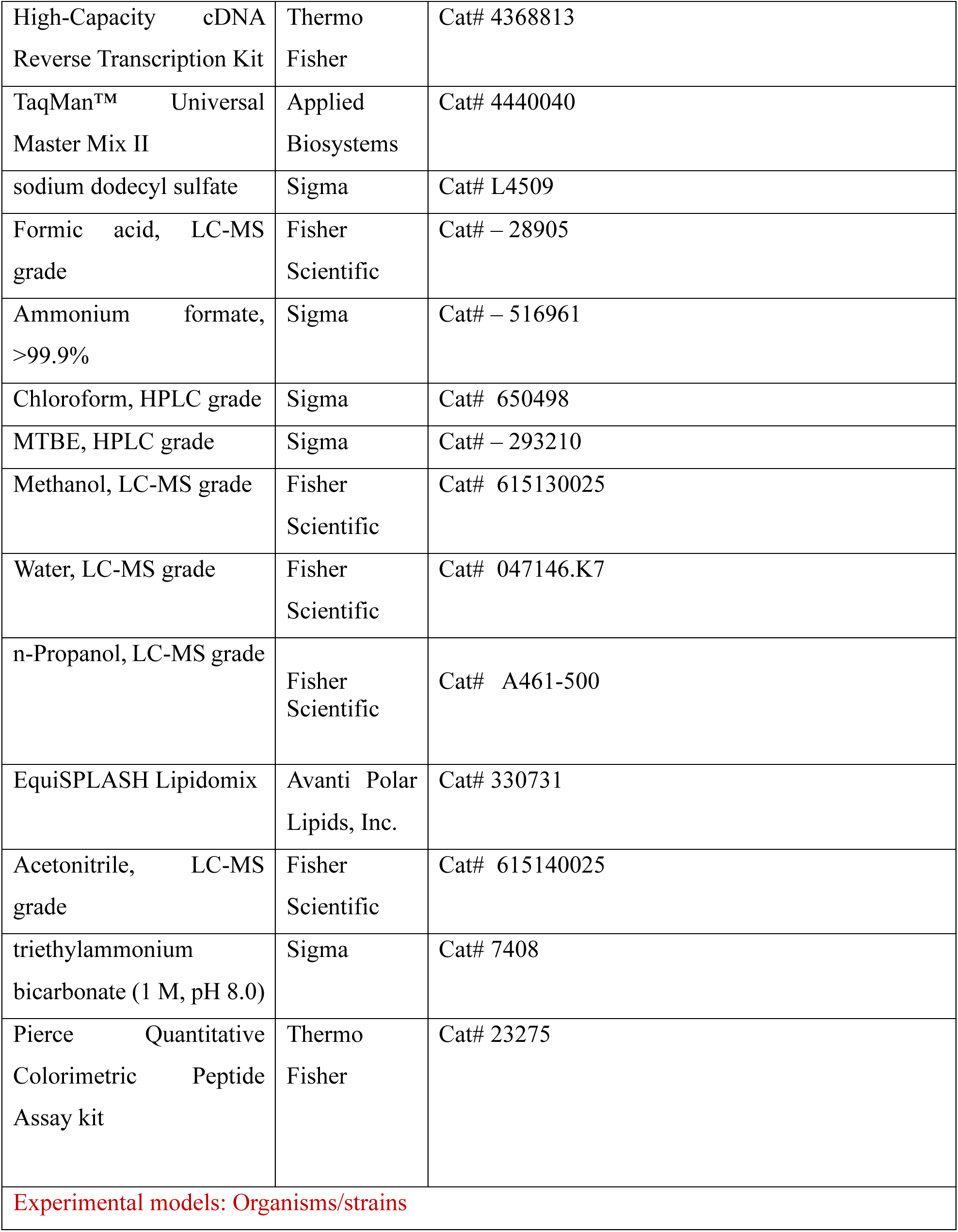
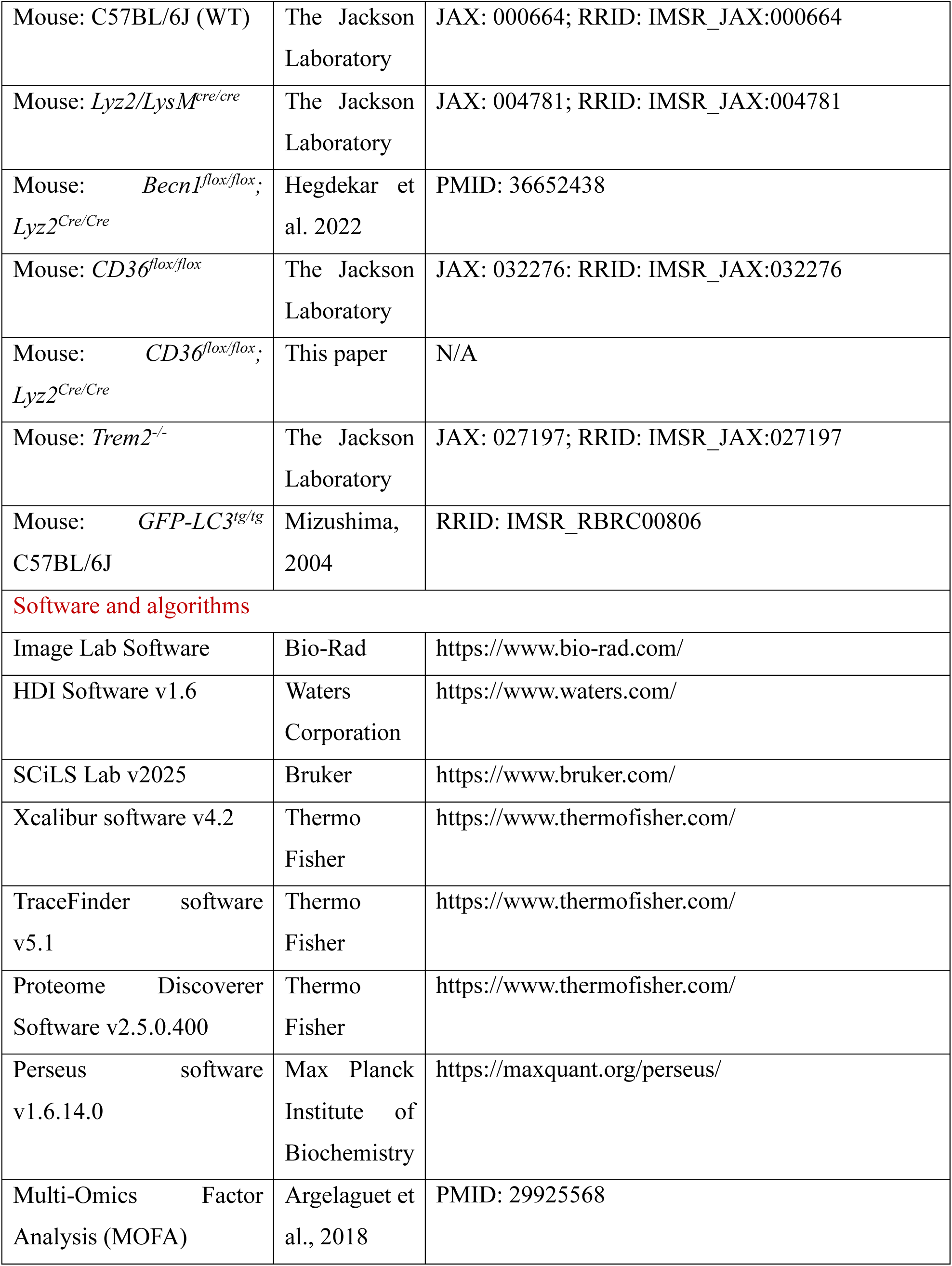
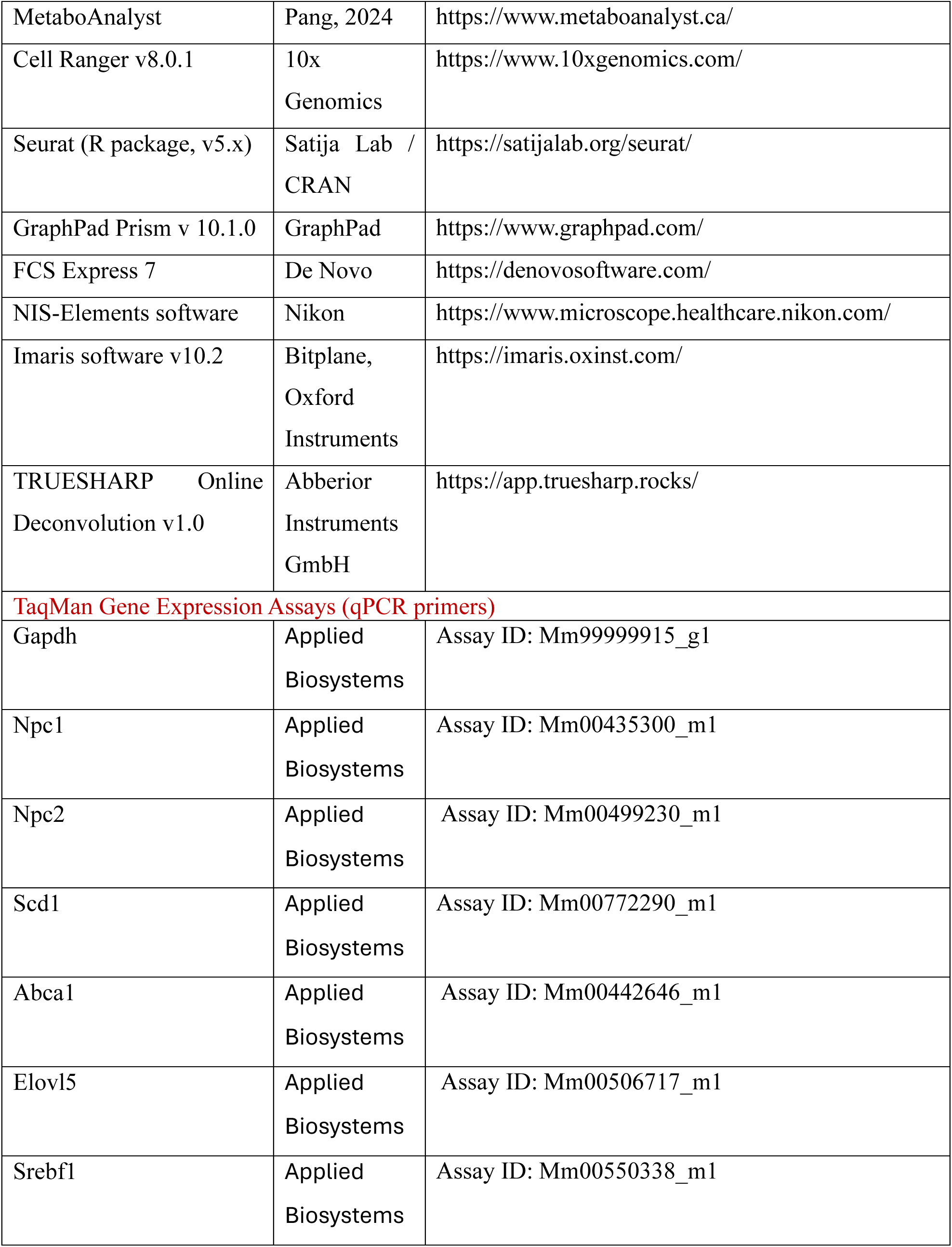
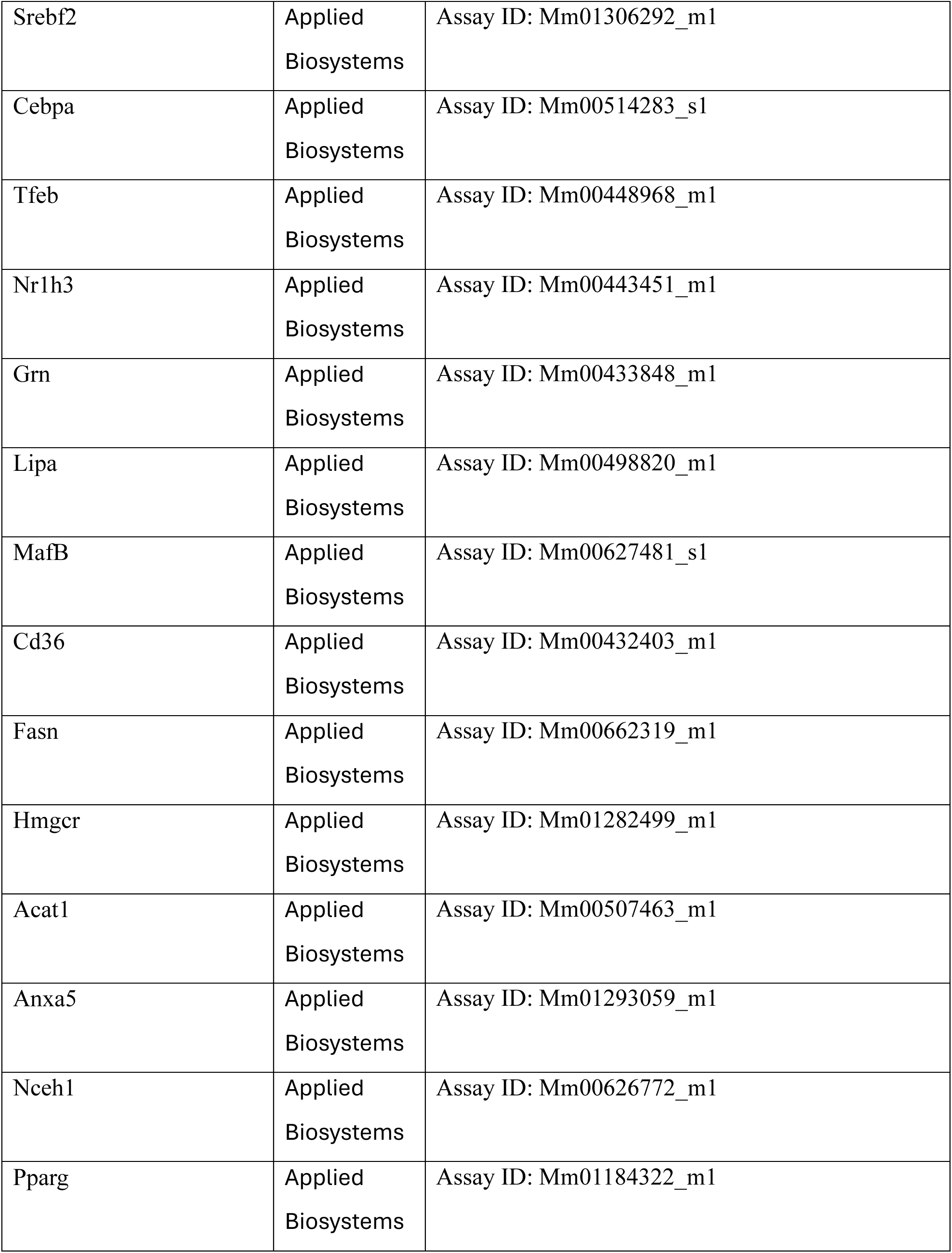
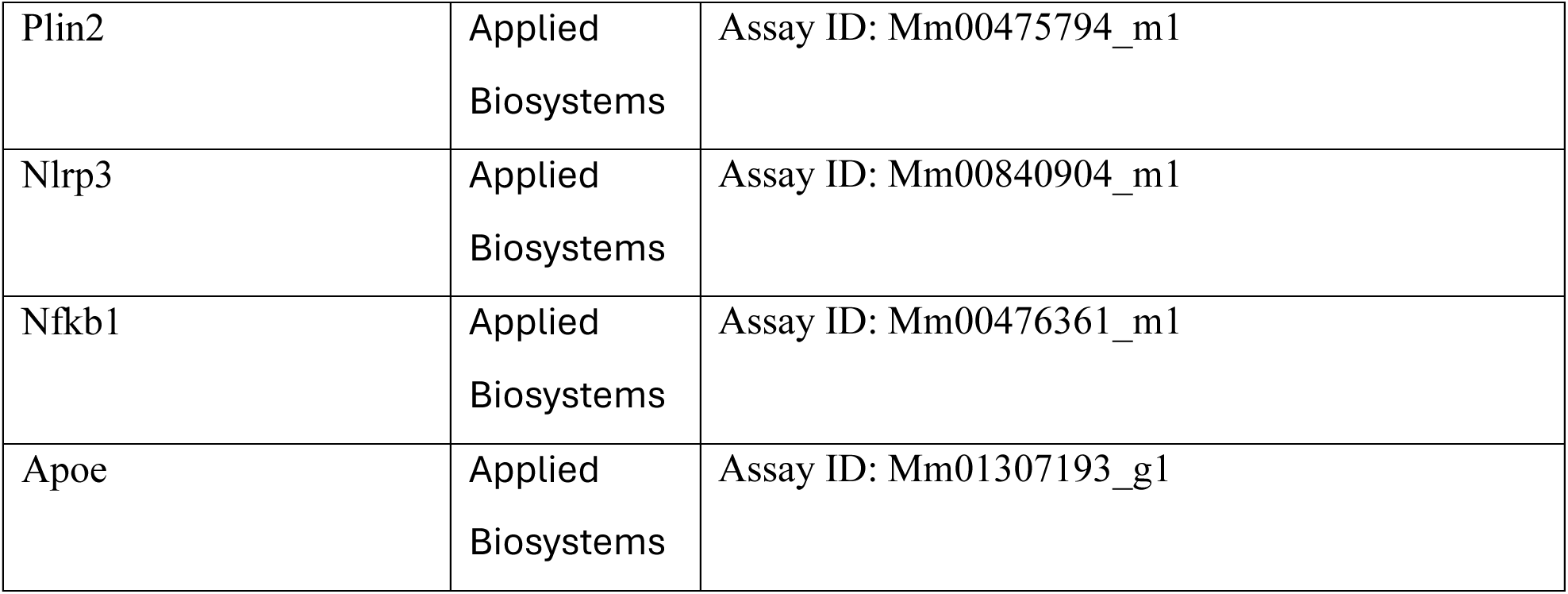
KEY RESOURCES TABLE

### Animals

All animal procedures were conducted in compliance with protocols approved by the University of Maryland’s Institutional Animal Care and Use Committee. All experimental mice were males on *C57Bl6/J* background. *Lyz2/LysM^cre/cre^*mice were crossed separately with *Becn^flox/flox^* and *Cd36^flox/flox^*mice to generate *Becn1^flox/flox^;Lyz2^Cre/Cre^* and *Cd36^flox/flox^;Lyz2^Cre/Cre^* monocyte and microglia-specific knockout mice. All offspring were genotyped for the presence of both the floxed and Cre alleles following the genotyping guidelines provided by The Jackson Laboratory. All mice were maintained on sterilized bedding in a specific pathogen-free (SPF) facility with a 12-hour light/dark cycle.

### Controlled Cortical Impact (CCI)

Traumatic brain injury (TBI) was induced in mice using a standard controlled cortical impact (CCI) device equipped with a microprocessor-regulated pneumatic impactor, as previously described^1,2^. Following anesthesia induction with 3% isoflurane, a 10-mm midline scalp incision was made to expose the skull. The skin and fascia were retracted, and a 4-mm craniotomy was performed over the central region of the left parietal bone. A moderate TBI was delivered using a 2.3-mm diameter impactor tip at a velocity of 4 m/s, deformation depth of 1.5 mm, and dwell time 100 ms. Sham-operated animals underwent the same surgical procedures, excluding the craniotomy and impact.

### DESI-MSI

Flash frozen brain tissues from four CCI and three sham mice were sectioned at 20 µm thickness using a cryostat (HM 550; Thermo Scientific, Waltham, MA). Tissues were dried in a desiccator for 15 minutes at room temperature prior to analysis. DESI-MSI was conducted using a HDMS select series Cyclic IMS mass spectrometer (Waters Corporation, Wilmslow, UK) equipped with a DESI XS source and a heated transfer line. The capillary voltage for the DESI XS source was set to 0.75 kV, with a cone voltage of 60 V, a source temperature of 150 °C, and a heated transfer line temperature of 300 °C for more efficient ion transfer. Samples were run at 50 µm spatial resolution, maintaining a scan speed of 1 second per scan during DESI-MSI analysis in positive ion mode. DESI flow was controlled by an Acquity Binary Solvent Manager (Waters Corporation, Milford, MA), maintaining a constant flow of 2 µL/min throughout the duration of the experiments. MSI data was first processed using High-Definition Imaging (HDI) software v1.6 (Waters Corporation, Milford, MA), where lipids were tentatively annotated based on accurate mass. Annotated data was then exported to SCiLS Lab v2025 (Bruker, Billerica, Massachusetts) for co-registration and visualization.

### Lipidomics

Lipid Extraction: Total lipid extracts from the lysosomes were prepared using methyl tert butyl ether (MTBE) lipid extraction protocol^3^ with slight modifications as described previously^4,5^. Briefly, 400 µL of cold methanol and 10 µL of internal standard (EquiSPLASH) were added to each sample. The sample was incubated at 4°C, 650 rpm shaking for 15 min. Next, 500 µL of cold MTBE was added followed by incubation at 4°C for 1 h with 650 rpm shaking. Cold water (500 µL) was added slowly, and the resulting extract was maintained 4°C, 650 rpm shaking for 15 min. Phase separation was completed by centrifugation at 8,000 g for 8 min at 4 °C. The upper, organic phase was removed and set aside on ice. The bottom, aqueous phase was re-extracted with 200 µL of MTBE followed by 15 min incubation at 4 °C with 650 rpm shaking. Phase separation was completed by centrifugation at 8,000 g for 8 min at 4 °C. The upper, organic phase was removed and combined with a previous organic extract. The latter was dried under a steady stream of nitrogen at 30 °C. The recovered lipids were reconstituted in 100 µL of acetonitrile:isopropanol:water (1:2:1, v/v/v).

Liquid Chromatography Tandem Mass Spectrometry (LC-MS/MS): Total lipid extracts were analyzed by liquid chromatography coupled to targeted tandem mass spectrometry (LC-MS/MS). The LC-MS/MS analyses were performed on an Ultimate 3000 Ultra High-Performance Liquid Chromatograph coupled to a Thermo TSQ Altis Tandem Quadrupole Mass Spectrometer (Thermo Scientific, San Jose, CA). LC-MS/MS methodology was adapted from the literature^6^ and previous publications^5,7^. The separation was achieved using an ACQUITY Amide BEH column (1.7 µm; 2.1 x 100 mm) column (Waters, Milford, MA) maintained at 45 °C. Mobile phase compositions for solvents A and B consisted of ACN/H_2_O (95:5, v/v) and (50:50, v/v) respectively, with 10 mM ammonium acetate The gradient profile had a flow rate of 0.6 mL min^−1^ and ramped from 0.1 to 20% B in 2 min, from 20 to 80% B in 3 min, dropped from 80 to 0.1% B in 0.1 min, and held 0.1% B for 2.9 min. Total chromatographic run time was 8.0 min. The injection volume was 2 μL. The auto-sampler was maintained at 7 °C. Electrospray ionization was achieved using either negative or positive mode. Mass spectrometry detection was done using selective reaction monitoring where predetermined precursor to product ion transitions were used. ESI source parameters were set as follows: voltage 3500 V in positive mode and −2500 V in negative mode, sheath gas (Arb) = 60, aux gas (Arb) = 15, sweep gas (Arb) = 1 and ion transfer tube temperature of 380 °C. Nitrogen was used as the nebulizer and argon as collision gas (1.5 mTor). The vaporizer temperature was set to 350 °C. LC-MS/MS data was acquired using Thermo’s Xcalibur software and data processing was achieved using Xcalibur 4.2 and TraceFinder 5.1. Additional data analysis was done using Prism 6 (GraphPad, La Jolla, CA) and MetaboAnalyst^8^. Lipidomics data has been uploaded to Mendeley data repository. Additional data is available upon request.

### BMDM isolation and culture and myelin phagocytosis

BMDMs were harvested from euthanized 12 – 14 weeks old female mice femurs and tibias under sterile conditions. Bones cleaned of muscle tissue and marrow were flushed using ice-cold PBS through a 23G needle into a collection tube. Cells were suspended by pipetting with a 1000 ul pipette, filtered through a 70 μm cell strainer to remove bone fragments and centrifuged at 400 × g for 5 minutes. Supernatant was discarded and pellet resuspended in RBC lysis buffer and left on ice for 10 minutes. Cells were centrifuged again (same speed and time) and the final white cells pellets resuspended and cultured in T75 flasks with BMDM media (RPMI containing 1% penicillin-streptomycin, 10% FBS, and 10% L929 conditioned media to provide M-CSF). Media was changed every 2-3 days. On day 7 cells were trypsinized and re-seeded on glass coverslips in 24 well plates at 2 ×10^5^ overnight, then treated with 100ul/ml purified mouse myelin^9,10^ 4 to 24 hours, as indicated. Cells were either fixated with 4% PFA for immunostaining or lysed with RIPA buffer for western blot^11,12^.

### Flow cytometry

Mice were anesthetized using isoflurane and perfused transcardially with cold saline to remove circulating blood. Cortical brain tissue was collected and mechanically dissociated using a 70 μm cell strainer. The resulting suspension was resuspended in RPMI-1640 medium. To facilitate enzymatic digestion, the suspension was treated with 10 U of papain, 10 mg/ml of DNase I, 1 mg/ml of collagenase-dispase, and 1 μl GolgiPlug containing brefeldin A, then incubated for 1 hour at 37°C on a shaker set to 200 rpm. Following digestion, leukocytes were isolated from brain tissue via a Percoll density gradient. Cells were first resuspended in 70% Percoll-HBSS and gently layered underneath a 30% Percoll-RPMI solution using a blunt-ended pipette tip. This gradient was centrifuged at 400 × g for 20 minutes without braking. The myelin layer was carefully removed from the top of the 30% interface, and immune cells were collected from the 30%/70% interphase and washed in RPMI medium to create single-cell suspensions. For flow cytometry, these brain leukocytes were washed in FACS buffer (1× HBSS containing 5% FBS, 0.1% penicillin-streptomycin, and sodium azide) and incubated with Fc Block diluted 1:50 for 10 minutes on ice to prevent nonspecific binding. Cells were then stained with fluorophore-conjugated antibodies against surface markers. After staining, they were fixed and permeabilized using BD Fixation/Permeabilization solution for 10 minutes, washed twice with BD Permeabilization/Wash Buffer, and incubated with intracellular antibody cocktails. Following a 30-minute incubation at 4°C, cells were washed again and resuspended in PBS for flow cytometry. Surface markers included CD45-eF450, CD11b-APCeF780 each used at 1:100 dilution. For viability staining, Zombie Aqua™ was prepared in DMSO and used at 1:50 dilution. Intracellular markers included TNF-PE-Cy7, 1:50, I-1β-PerCP-eF710, 1:100, LC3B-FITC 1:100, and SQSTM1/p62-AF647, 1:50.^13^ Flow cytometric data were acquired using a 5-laser Cytek Aurora cytometer with SpectroFlo 3.30 software. Unmixing was done using single color controls (SCC). SCCs for dyes (BODIPY, FluoroMyelin, and Zombie) were made by cells and SCCs for antibodies made by compensation beads. Analysis was done with FCS Express™ 7.

### CD11B^+^ cell isolation for single cell RNA sequencing and FACS

Cells were isolated using Adult Brain Dissociation Kit (ABDK) following manufacturer’s instructions, with modifications.^14,15^ All procedures were performed on ice/4 °C, as indicated; all plasticware was pre-coated with coating buffer (3% BSA in DPBS-CMF) to prevent cell adhesion. Anesthetized mice were transcardially perfused with 20 mL cold perfusion solution (30 mL DPBS-CMF with 2 U/mL heparin). Brains were dissected on ice and regions of interest micro-dissected into petri dishes containing 1 mL cold Enzyme Mix 1. 30 µL of Enzyme Mix 2 was added to each C Tube containing tissue, attach tubes to gentle MACS Octo Dissociator, and run the customized protocol.^15^ After dissociation, cells were filtered through 70 µm mesh into 50 mL tubes pre-wet with 500 µL cold DPBS-CMF. C Tubes were rised with 9.5 mL DPBS-CMF and add to the filters then centrifuged at 300 × g for 8 min at 4 °C. Supernatant aspirated carefully. For debris removal, the pellet was gently resuspend in 1550 µL DPBS-CMF, mixed with 450 µL cold Debris Removal Solution, and overlaid with 2 × 1 mL DPBS-CMF. The mix was centrifuged at 3000 × g, 10 min, 4 °C then top two layers were discarded. The remnant was diluted to 5 mL with cold DPBS-CMF, inverted gently 5 times and centrifuged at 1000 × g for 10 min. Supernatant was aspirated and the pellet was resuspend in 90 µL MACS buffer [95 ml DPBS-CMF with 5 ml FBS (5% final) and 200 µL EDTA 0.5M (final 1mM)]. 10 µL CD11b microbeads was added and mixed by pipetting gently and left to incubate on ice for 15 min. Then 1 mL MACS buffer added and mixed by pipetting and centrifuged at 300 × g for 8 min at 4 °C. Meanwhile LS columns set on the magnetic stand and washed suing 2 ml MACS buffer. Also, 30 µm filters were pre-wet by using 1 ml MACS buffer. Once centrifuge was done, supernatant was aspirated and the pellet was resuspended with 500 MACS buffer and loaded to LS the column and rinsed with 2 mL MACS buffer (washing column with 3 mL MACS buffer is recommended). CD11b+ cells were eluted by removing columns from the magnet and flushing with 2 × 5 mL MACS buffer. Eluates spinned at 300 × g for 8 min, aspirated carefully (pellet may be invisible). For downstream applications like scRNA-seq or FACS, pellet resuspended in 400 µL cold 1× PBS and original tube was rinsed with 800 µL PBS then combined and spinned again, and finally resuspended in MACS buffer (for scRNAseq) or FACS buffer (for FACS with that was done using BD Aria sorter).

### Single cell RNA sequencing and Seurat Analysis

Single cells in MASC buffer were subjected to encapsulation and library preparation that was carried out using the 10x Genomics Chromium v2 platform, following the manufacturer’s protocols. Each sample was processed as an independent batch, with a capture target of at least10,000 cells. Both cDNA and library amplifications were performed with 12 PCR cycles. Sequencing was conducted on the Illumina NovaSeq 6000 S2 flow cell (RRID:SCR_016387), according to the manufacturer’s recommendations. An initial low-depth sequencing run (∼1,000 reads per cell) was performed on an Illumina NextSeq instrument to validate input concentration and sample quality. Final sequencing was performed at approximately 100,000 reads per cell using the Illumina NovaSeq platform. Post-sequencing, library complexity and saturation were assessed using the Preseq package. Samples with saturation levels below 60% were sequenced further to ensure sufficient coverage, guided by Preseq projections. All samples underwent alignment and preprocessing using the Cell Ranger pipeline (version: cellranger-8.0.1, Transcriptome: GRCm39-2024-A). Analysis was done using Seurat (R package, v5.x). Quality control filtering was applied to retain cells that met the following criteria: detection of 200 to 10,000 genes per cell, total unique molecular identifier (UMI) counts between 1,000 and 15,000, and no more than 20% of reads aligning to mitochondrial genes resulting an average of 15,371 cells from sham samples and 12,493 cell from CCI samples. The majority of sn/scRNAseq analyses are comparisons between relatively stable cell populations, such as cells from organisms of different genotypes, ages or established disease states. Consistent with these dynamic changes, we detected significantly higher numbers of different mRNA transcripts in cells from injured as compared to sham mice (average read depth of 4,662/cell for sham, 10,289/cell for CCI; Supp. Figure S2A). To account for this, we performed stringent quality control assessments including number of RNA features, mitochondrial RNA content, normality of distribution^16^. The higher number of RNA features in TBI was justified by cell cycle analysis (Supp. Figure S2B, Figure 2F) indicating proliferation of monocytes after TBI. Individual samples as Seurat objects were integrated using IntegrateLayers() Seurat function. Differentially expressed (DE) genes were defined by FindAllMarkers() Seurat function that uses Wilcoxon rand sum test. Identity assignment to Seurat clusters done manually using canonical microglia and macrophages markers. These included canonical homeostatic markers (e.g. *Tmem119*)^17^ which were used to identify homeostatic microglia clusters (Hom_MG_1-5). Additional clusters expressing high levels of homeostatic markers like Tmem119 but also positive for some activation markers (e.g. *Lpl* and *Cybb*) were designated as surveillance microglia (Surv_MG_1-5). Clusters with high expression of disease associated microglia (DAM) markers (e.g. *Lyz2*)^18^ along with high microglial markers were identified as DAM_1-5. Clusters with high DAM markers and lower expression of microglial markers were identified as brain associated macrophages (BAM_1-7). BAM identity was further confirmed based on expression of infiltration (e.g. *Ccr2*) or macrophage differentiation (e.g. *Pf4*) markers^19^. Additional minor clusters included immune cells such as dendritic cells, neutrophils, NK and T cells, as well as astrocytes and mature and immature oligodendrocytes (Supp. Figure 2C).

### Western blot analysis

Cell lysates were obtained by directly lysing BMDMs cultured in 24-well plates using RIPA buffer. Proteins from cellular lysates were separated on 4–20% gradient SDS-PAGE gels, followed by transfer onto PVDF membranes using a semi-dry blotting apparatus (Bio-Rad). The membranes were first blocked with 5% non-fat milk in TBST buffer (Tris-buffered saline containing 0.05% Tween 20), then incubated overnight at 4°C with primary antibodies diluted in 1% BSA in TBST (β-Actin/ACTB,1:10,000; SQSTM1/p62,1:1000; LC3,1:1000). The next day, membranes were treated with HRP-conjugated secondary antibodies for 1 hour at room temperature in blocking solution. Protein signals were detected using either SuperSignal West Dura or SuperSignal West Pico chemiluminescent substrates and visualized with the Chemi-Doc imaging system (Bio-Rad, Universal Hood II). Band intensities were quantified using Image Lab software (Bio-Rad) and normalized to loading controls.

### Immunofluorescence staining

Mice were anesthetized using isoflurane and perfused transcardially with cold saline, followed by 4% paraformaldehyde (PFA, pH 7.4). Dissected brains were post-fixed in 4% PFA and cryoprotected in 30% sucrose before being sectioned into 20-μm-thick frozen slices, following previously established procedures^1^. For each immunofluorescence experiment, four coronal sections spaced 1–1.2 mm apart across the lesion site were collected per animal. Tissue sections were blocked with 5% goat serum in 1× phosphate-buffered saline (PBS) containing 0.025% Triton X100. For lysosomal labeling, sections were first treated with 0.1 M glycine in PBS for 30 minutes, followed by permeabilization with 0.2% saponin in PBS for another 30 minutes. They were then blocked in PBS containing 0.04% saponin, 5% goat or donkey serum, and 0.05% BSA. Sections were incubated overnight at 4°C with primary antibodies, followed by a 2-hour incubation at room temperature with appropriate secondary antibodies in blocking solution. DAPI was used to counterstain nuclei. BODIPY staining was done as a separate step right after permeabilization. BODIPY stock was made by reconstitution of the BODIPY™ 493/503 in DMSO (1 mg/ml). Stock was diluted 1:5000 in PBS and used applied on the sections for 1 hour in dark. Then washed three times with PBS and blocked.

### Epifluorescence image acquisition and quantification

Fluorescence images were captured using a Nikon Eclipse Ti-E/Ni-E microscope and analyzed with Nikon Elements software (version 4.12.01). Nikon 20X/0.75 Plan-Apochromat Lambda DIC objective lens (cellular neutral lipid and lipid droplet accumulation) or 60X /1.40 Plan-Apochromat Lambda DIC objective lens (BODIPY engulfment inside lysosome). Emission wavelengths used for detection included 460 nm for DAPI, 535 nm for BODIPY and Alexa Fluor 488, 620 nm for Alexa Fluor 546 and Alexa Fluor 568, 670 nm for Alexa Fluor 633, and 756 nm for Alexa Fluor 750. Z-stacks were captured at 1 μm intervals and processed with the NIS-Elements Extended Depth of Focus (EDF) algorithm to reconstruct a single in-focus composite image. Consistent exposure times were maintained across all tissue sections within each experimental group. Quantification was carried out using Elements software. Nuclei were identified using the Spot Detection algorithm, while immunofluorescent marker-positive cells were detected using the Detect Regional Maxima algorithm in combination with global thresholding, as previously described^2^. The number of marker-positive cells was normalized to the total number of nuclei per image.

### Confocal image acquisition and 3D rendering

Confocal fluorescence images were acquired with the Nikon CSU-W1 microscope equipped with 405, 488, 561, and 647 lasers, using a 60x (1.49 NA) TIRF oil-immersion objective and Nikon Elements software. For three-dimensional rendering, confocal z stacks were taken with a 60x objective equipped with a 1.5x magnifying lens, to achieve a magnification of 102x and a z-step size of 0.2um. Images were denoised and deconvolved using automatic deconvolution algorithms in Nikon Elements and were reconstructed in three dimensions in Imaris Bitplane software Individual microglial surface of fluorescence images were reconstructed using the ‘surface’ feature and default parameters to create a volumetric boundary of the cell. Following all the lysosomal signal outside of the reconstructed surfaces were masked to ‘0’, retaining only lysosomal signal inside the reconstructed surfaces. After masking, both lysosomal surfaces and BODIPY-positive signals were reconstructed using the ‘surface’ feature and ‘machine learning’ algorithm parameters. The two surfaces (BODIPY-positive and lysosomes) were merged to create the final rendering. Representative images were deconvolved using TRUESHARP online deconvolution (Abberior). All analysis were performed on raw images.

### Lysosomal enzyme assay

The enzymatic activities of lysosomal proteins cathepsin D (CTSD) and alpha-N-acetylglucosaminidase (NAGLU) were measured using respective fluorometric assay kits following the manufacturers’ protocols. Enzyme activity levels in lysosomal fractions and total cell or tissue lysates were quantified based on changes in absorbance or fluorescence per microgram of protein. To assess cytosolic enzyme activity, values were calculated as a proportion of the total enzyme activity in the whole cell or tissue lysate, normalized per microgram of total protein.

### Preparation of lysosome enriched fraction

Lysosomal fractions were isolated from cortical tissue of both sham and injured mice using previously established protocols^1^ using the Lysosomal Enrichment Kit for Tissue and Cultured Cells following the manufacturer’s recommended procedure.

### Lysosomal proteomics

We collected lysosomal samples and stored them at -80 °C until assay. Cell lysis and protein digestion were performed similar as previously described^20,21^. samples were lysed in a lysis buffer containing 5% sodium dodecyl sulfate, 50 mM triethylammonium bicarbonate (1 M, pH 8.0). Proteins were extracted and digested using S-trap micro columns (ProtiFi, NY). The eluted peptides from S-trap column were dried, and peptide concentration was determined using Pierce Quantitative Colorimetric Peptide Assay kit, after reconstituted in 0.1% formic acid. LC-MS/MS-based proteomic analysis was conducted on a nanoACQUITY Ultra-Performance Liquid Chromatography system (Waters Corporation, Milford, MA USA) coupled to an Orbitrap Fusion Lumos Tribrid mass spectrometer (Thermo Scientific, San Jose, CA USA) similar to our previous work^20,22,23^. Peptide separation was effected on a nanoACQUITY Ultra-Performance Liquid Chromatography (UPLC) analytical column (BEH130 C18, 1.7 µm, 75 µm x 200 mm; Waters Corporation, Milford, MA, USA) using a 185-min linear gradient with 3-40% acetonitrile and 0.1% formic acid. Mass spectrometry conditions were as follows: Full MS scan resolution of 240,000, precursor ions fragmentation by high-energy collisional dissociation of 35%, and a maximum cycle time of 3 seconds. The Pierce HeLa Protein Digest Standard was injected between runs as an instrument quality control to monitor system performance. The resulting mass spectra were processed using Thermo Proteome Discoverer (PD, version 2.5.0.400, Thermo Fisher Scientific) and searched against a UniProt mouse (Mus musculus) reference proteome (release 2022.06, 17180 entries) using Sequest HT algorithm. Search parameters include carbamidomethylation of cysteines (+57.021 Da) as a static modification, methionine oxidation (+15.995 Da) as a dynamic modification, precursor mass tolerance of 20 ppm, fragment mass tolerance of 0.5 Da, and trypsin as a digestion enzyme. Tryptic missed cleavages were restricted to a maximum of two, with peptide lengths set between 6 and 144 residues. For protein quantification, the Minora feature detector, integrated in the PD, was used as described previously^24^. To ensure high data quality, proteins were further filtered to a 1% false discovery rate (FDR) threshold, calculated with the Percolator algorithm. Next, protein abundance values exported from PD were post-processed using Perseus software. Proteins with missing values were excluded to improve data quality. The quantitative protein data were log_2_ transformed and further normalized using median centering. Differentially expressed proteins (DEPs) were identified using a two-tailed Student’s t-test (adjusted p-value < 0.05). Ingenuity Pathway Analysis (IPA) software (Qiagen, Germantown, MD) was used to identify dysregulated pathways and biological processes. Metabolanalyst (version 5.0) was used to generate PCA plots and heatmaps. The mass spectrometry proteomics data have been deposited to the ProteomeXchange Consortium via the PRIDE partner repository with the project accession number PXD067157.

### Quantitative real time PCR

Total RNA was extracted from cortex using TRIzol reagent and RNeasy Mini Kit (QIAGEN) following the manufacturer’s protocol. The RNA was reverse transcribed into complementary DNA (cDNA) using the High-Capacity cDNA Synthesis Kit according to the manufacturer’s instructions. Quantitative real-time PCR (qRT-PCR) was carried out using the TaqMan Universal Master Mix II. For each reaction, 2× Master Mix, 1 μl of cDNA (equivalent to 50 ng of input RNA), and the appropriate TaqMan Gene Expression Assay were combined in a final volume of 20 μl. All reactions were performed in duplicate. TaqMan Gene Expression Assays (primers) targeting mouse transcripts were utilized (detailed informations of each listed in the resources table). PCR amplification and data collection were conducted on a 7900HT Fast Real-Time PCR System (Applied Biosystems) using the manufacturer’s software. The thermal cycling conditions included an initial hold at 50 °C for 2 minutes and 95 °C for 10 minutes, followed by 40 cycles of denaturation at 95 °C for 15 seconds and annealing/extension at 60 °C for 1 minute. Gene expression levels were normalized to Gapdh, and relative expression was calculated accordingly.

